# Upregulation of the lncRNA *XACT* sustains pluripotency, blocks lineage specification, and drives germ cell tumor-like transcriptional programs in human pluripotent stem cells

**DOI:** 10.1101/2025.10.02.679492

**Authors:** Kenji Izumi, Nami Motosugi, Akiko Sugiyama, Natsumi Kurosaki, Yumi Iida, Misaki Higashiseto, Keiko Yokoyama, Ayumi Sasaki, Atsushi Izumi, Tohru Kiumra, Mitsutoshi Yamada, Akihiro Umezawa, Hidenori Akutsu, Hitoshi Ishimoto, Natsuhiko Kumasaka, Atsushi Fukuda

**Affiliations:** Department of Molecular Life Sciences, Division of Basic Medical Science and Molecular Medicine, Tokai University School of Medicine, Isehara, Kanagawa, Japan; Department of Obstetrics and Gynecology, Tokai University School of Medicine, Isehara, Japan; Department of Life Science Support, Research Innovation Center, University Hospitals Sector, Tokai University, Isehara, Kanagawa, Japan; Laboratory of Stem Cell Biology, Department of Biosciences, Kitasato University School of Science, Kanagawa, Japan; Department of Obstetrics and Gynecology, Keio University School of Medicine, Tokyo, Japan; Center for Regenerative Medicine, National Center for Child Health and Development, Tokyo, Japan; The Institute of Medical Sciences, University of Tokyo, Tokyo, Japan; The Institute of Medical Sciences, Tokai University, Isehara, Kanagawa, Japan; Micro/Nano Technology Center, Tokai University, Hiratsuka, Kanagawa, Japan

## Abstract

Long noncoding RNAs (lncRNAs) represent a vast class of regulatory transcripts and are spatiotemporally controlled, yet only a few have been functionally implicated in human development. Here, we identify the X-linked lncRNA *XACT*, abundantly but transiently expressed during early human embryogenesis, as a critical regulator of pluripotency, lineage specification, and cancer-like states. In human pluripotent stem cells (hPSCs), *XACT* overexpression—but not depletion—sustains self-renewal without exogenous factors and prevents lineage commitment. Mechanistically, *XACT* upregulation drives hyper-elevation of the core pluripotency factors OCT4 and NANOG at the protein level by repressing their 3′ untranslated regions (UTRs). *XACT* overexpression confers context-dependent states: in standard hPSC medium it promotes a naïve-like program, whereas in the absence of exogenous factors it drives transcriptomic states resembling testicular germ cell tumors, linking misregulation of a developmentally restricted lncRNA to tumorigenic potential. In hPSC-based models of post-implantation development, *XACT* expression normally declines, whereas its sustained expression disrupts embryonic progression, while depletion has little effect. Finally, transcriptomic analysis of post-implantation human embryos showed that *XACT* levels correlate positively with pluripotency-associated gene networks. Together, these findings establish *XACT* as a potent, human-specific modulator of pluripotency and early embryogenesis, and suggest that its aberrant upregulation may underlie both developmental failure and germ cell tumorigenesis.

## Introduction

Early human development is governed by species-specific transcriptional programs, including dynamic regulation of transcription factors and noncoding RNAs^1–3^. Many genes are transiently expressed during pre-implantation stages, coinciding with the onset of lineage commitment^3–5^. Functional interrogation of such regulators—through both loss- and gain-of-function approaches—is essential for understanding human-specific developmental mechanisms. However, reverse genetic studies in human embryos are severely constrained by sample availability and ethical considerations, leaving critical determinants of human embryogenesis largely undefined.

Human pluripotent stem cells (hPSCs) and their naïve counterparts, which capture properties o F epiblast and morula-stage cells, respectively, provide a tractable platform to model early development^6–8^. Around the morula stage, zygotic genome activation occurs, and the core pluripotency factors OCT4, NANOG, and SOX2 are induced prior to lineage specification^3,9,10^. While these factors are transiently expressed in early embryos and widely used as markers of pluripotency—including naïve states—similar transcriptional signatures are also observed in cancers^11,12^. Dissecting how such transcriptional states are established and maintained in hPSCs is therefore critical both for developmental biology and for clinical applications.

In parallel with transcription factors, numerous long noncoding RNAs (lncRNAs) are expressed during pre-implantation development^3,13–15^. Although many transcription factors directing developmental transitions have been defined, only a handful of lncRNAs have been implicated as pivotal regulators. One such candidate is *XACT*, an X-linked lncRNA uniquely abundant during human pre-implantation stages and robustly expressed in hPSCs ^15–17^. We previously reported that *XACT* depletion does not impair pluripotency but accelerates neuronal differentiation^16^, implicating it in lineage control. Notably, *XACT* expression is normally silenced upon hPSC differentiation^18^, suggesting that its sustained expression could perturb developmental trajectories.

Here, we combined loss- and gain-of-function approaches in hPSCs to dissect the role of *XACT*. Strikingly, *XACT* overexpression—but not repression—profoundly altered differentiation, early embryonic modeling, and molecular states. Elevated *XACT* drove hyper-elecation of OCT4 and NANOG proteins by repressing their 3’ untranslated regions (UTRs), sustaining self-renewal without exogenous supplements and blocking lineage specification. Transcriptomic analyses further revealed that the consequences of *XACT* overexpression were strongly dependent on culture context: under conventional hPSC conditions it reinforced a naïve-like pluripotent program, whereas in the absence of exogenous factors it redirected cells toward transcriptional states closely resembling testicular germ cell tumors. In hPSC-based models of post-implantation development, sustained *XACT* expression disrupted embryonic progression. Consistently, single-cell transcriptomic data from human embryos showed that *XACT* expression correlates with pluripotent cell populations.

Together, these findings identify *XACT* as a potent regulator of pluripotency acting through post-transcriptional modulation of core transcription factors, and suggest that misregulation of a single lncRNA can derail human development and promote tumor-like states.

## Results

### *XACT* upregulation causes hyper-elevated OCT4 and NANOG protein expression across pluripotent and undirected differentiation contexts

To elucidate the potential role of *XACT* RNA during early human differentiation, we employed both loss- and gain-of-function strategies in male iPSCs (*XACT*^WT/Y^). Loss-of-function was achieved using previously established *XACT* knockout lines (*XACT*^Δ/Y^-1, −11)^16^, while gain-of-function was performed using a doxycycline (Dox)-inducible CRISPR activation (CRISPRa) system based on PiggyBac-mediated expression of dCas9-VPR and guide RNAs targeting the *XACT* locus (*XACT*^VPR^-3,-6) (Fig. 1a: schematic and Supplementary Fig. S1a)^16,18^.

**Figure 1.**
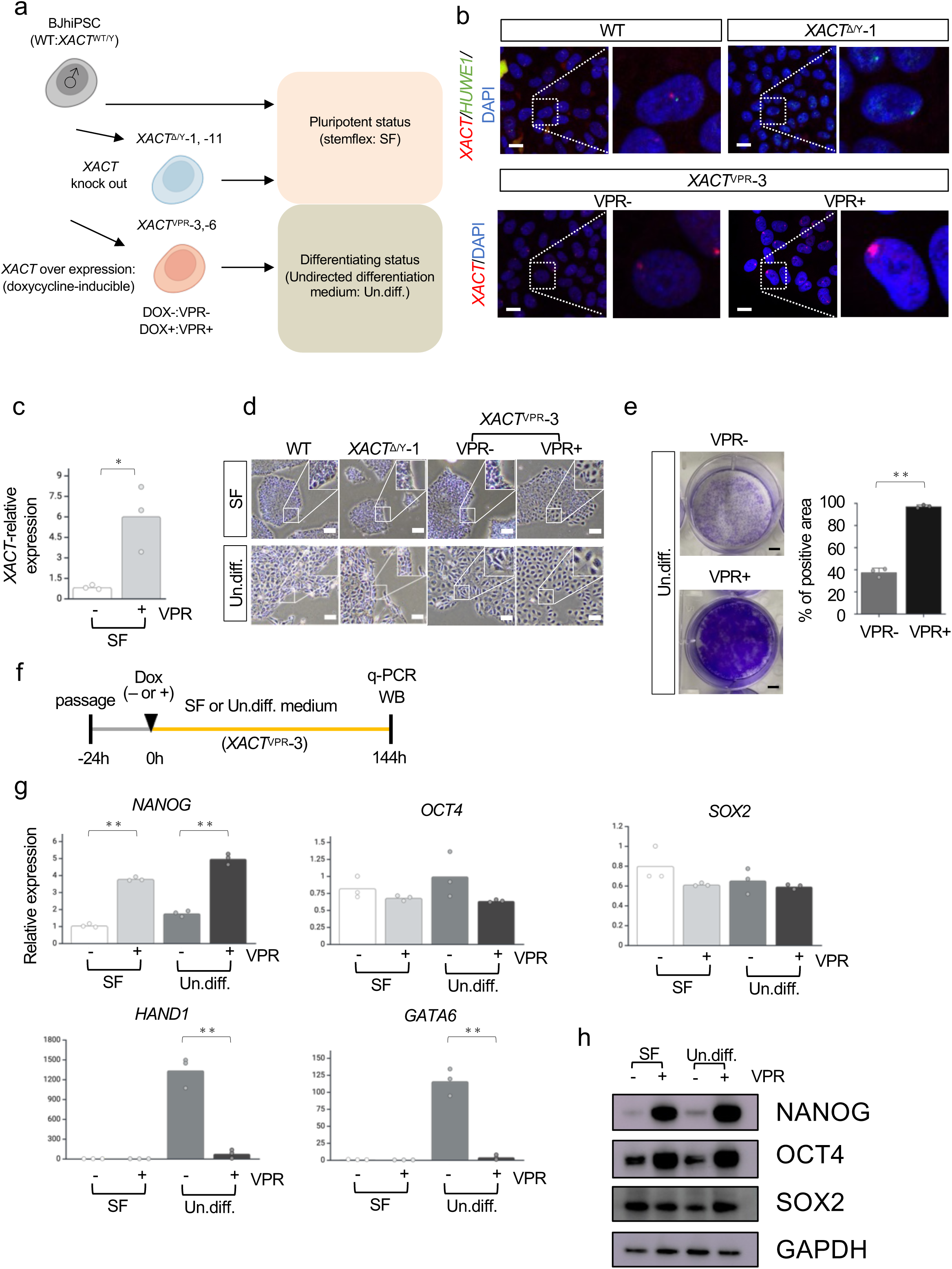
*XACT* upregulation induces hyper-elevation of NANOG and OCT4 protein levels. **(a)** Schematic overview of the bi-directional functional approach to investigate the role of *XACT* in pluripotency and differentiation. Loss-of-function was assessed using two independent *XACT* knockout lines (*XACT*^Δ/Y^-1 and *XACT*^Δ/Y^-11), while gain-of-function was achieved by CRISPR activation with a doxycycline (Dox)-inducible dCas9:VPR system (two independent lines: *XACT*^VPR^-3 and *XACT*^VPR^-6). Data for *XACT*^VPR^-6 are shown in Supplementary Figures. Dox-treated *XACT*^VPR^ cells are indicated as VPR+. Cells were maintained in StemFlex (SF) medium under pluripotent conditions or in DMEM/F12 basal medium lacking bFGF/Activin for undirected differentiation (Un.diff.). (b) RNA-FISH analysis of *XACT* in *XACT*^WT/Y^, *XACT*^Δ/Y^-1, and *XACT*^VPR^-3 cells. HUWE1 was used as a control probe. In *XACT*^VPR^ cells, Dox was added for 72 h prior to RNA-FISH. Scale bar, 20 µm. (c) qPCR analysis of *XACT* expression in *XACT*^VPR^ cells upon Dox induction. Each dot represents a biological replicate. *P < 0.05 (Student’s t-test). (d) Bright-field images of WT, *XACT*^Δ/Y^-1, and *XACT*^VPR^-3 cells cultured in SF or Un.diff. conditions. Scale bars, 100 µm. Data for *XACT*^Δ/Y^-11 and *XACT*^VPR^-6 are shown in Supplementary Fig. S1d. (e) Crystal violet assay of cell growth in Un.diff. conditions. Cells were seeded at 1 × 10^5^ and stained at 120h post-Dox. Each dot represents a biological replicate; bar graphs indicate mean values. *P < 0.05; **P < 0.01 (Student’s t-test). (f) Experimental design for mRNA and protein expression analysis of pluripotency (NANOG, OCT4, SOX2) and differentiation (HAND1, GATA6) markers in *XACT*^VPR^-3 cells under SF or Un.diff. conditions. Protein expression was examined by western blotting (WB). Data for *XACT*^VPR^-6 are shown in Supplementary Fig. S1. (g) qPCR analysis. *P < 0.05; **P < 0.01 (Student’s t-test). (h) WB analysis of core pluripotency factors. GAPDH served as a loading control. Representative images are shown (uncropped blots and biological replicates are in Supplementary Fig. S10).

Consistent with previous reports^16^, RNA fluorescence in situ hybridization (RNA-FISH) confirmed the absence of *XACT* RNA in *XACT*^Δ/Y^-1, −11 cells (Fig. 1b; Supplementary Fig. S1b), whereas *XACT*^VPR^ lines exhibited strong *XACT* RNA signals following Dox treatment (hereafter *XACT*^VPR+^) at 144 hours post-induction (Fig. 1b and Supplementary Fig. S1b). Quantitative PCR (qPCR) analysis showed significantly upregulation (>3.4-fold up) of *XACT* in *XACT*^VPR+^ cells compared to *XACT*^VPR-^ controls at 144 hours after induction (Fig. 1c and Supplementary Fig. S1c), confirming the efficacy of the CRISPRa system.

In standard hPSC culture (StemFlex medium, SF)^19^, *XACT XACT*^VPR+^ cells developed less densely packed colonies after 144 hours of Dox treatment, while *XACT*^Δ/Y^ cells retained typical morphology without notable alterations (Fig. 1d and Supplementary Fig. S1d), consistent with prior reports^16^. This suggests that *XACT* activation, rather than repression, can influence cellular phenotype.

To assess whether *XACT* modulation affects differentiation, cells were cultured in a spontaneous differentiation medium lacking bFGF and Activin (Undirected differentiation: Un.diff.; Fig. 1a), which are essential for pluripotency maintenance^20,21^. After 144 hours under Un.diff. conditions, typical hPSC morphology was lost in all cells (Fig. 1d and Supplementary Fig. S1c). Strikingly, *XACT*^VPR+^ cells maintained colony morphology similar to SF conditions (Fig. 1d and Supplementary Fig. S1c) and exhibited significantly enhanced proliferation 120 hours after induction compared to *XACT*^VPR-^ controls (Fig. 1e and Supplementary Fig. S1e). Together with previous data showing that *XACT* depletion does not prevent differentiation^16^, these results indicate that *XACT* upregulation, rather than loss, can influence pluripotent status.

We next examined core pluripotency gene expression (NANOG, OCT4, and SOX2) and differentiation markers (HAND1 and GATA6)^18^ by qPCR and protein expression by western blotting under both SF and Un.diff. conditions (Fig. 1f). In SF medium, NANOG was significantly upregulated (>3 fold up) in *XACT*^VPR+^ cells compared to *XACT*^VPR-^ cells (Fig. 1g and Supplementary Fig. S1f), whereas OCT4 and SOX2 mRNA levels remained unchanged (Fig. 1g and Supplementary Fig. S1f).

Under Un.diff. conditions, differentiation-associated genes GATA6 and HAND1 were strongly induced (>17.4 fold up) in *XACT*^VPR-^ cells, but this induction was largely suppressed in *XACT*^VPR+^ cells (Fig. 1g and Supplementary Fig. S1f). NANOG mRNA was also significantly upregulated and SOX2 tended to be upregulated, while *OCT4* mRNA remained comparable to controls (Fig. 1g and Supplementary Fig. S1f). These results indicate that *XACT* upregulation fails to activate differentiation related genes with maintenance of core pluripotency factor expression in the absence of key extrinsic factors such as bFGF and Activin.

We then assessed protein expression of core pluripotency factors. NANOG protein was hyper-elevated in *XACT*^VPR+^ cells under both SF and Un.diff. conditions (Fig. 1h and Supplementary Fig. S1g). Notably, OCT4 protein was also markedly upregulated in *XACT*^VPR+^ cells under both conditions (Fig. 1h and Supplementary Fig. S1g), despite unchanged mRNA levels, suggesting that *XACT* may enhance OCT4 protein via post-transcriptional mechanisms, such as mRNA stability, translation efficiency, or protein stability. Regarding to SOX2, the protein level did not change in SF condition, but it showed slight increases under Un.diff., consistent with qPCR results (Fig. 1h and Supplementary Fig. S1g).

Taken together, these findings demonstrate that *XACT* upregulation prevents activation of differentiation related genes while the effects on OCT4 and NANOG protein levels are particularly pronounced and largely independent of culture conditions.

### *XACT* upregulation induces heterogeneous NANOG and OCT4 expression with propagation and reversibility

To examine core pluripotency factor protein expression in *XACT*^VPR+^ cells at single-cell resolution, we performed immunofluorescence (IF) analysis for NANOG, OCT4, and SOX2 in both SF and Un.diff. conditions (Fig. 2a). Consistent with western blot results (Fig. 1h), NANOG and OCT4 signals were markedly stronger in *XACT*^VPR+^ cells than in *XACT*^VPR-^ controls in both SF and Un.diff. conditions while SOX2 expression levels were not greatly induced in *XACT* upregulation (Fig. 2b and Supplementary Fig. 2a). Notably, NANOG and OCT4 exhibited pronounced heterogeneous expression across the *XACT*^VPR+^ population (Fig. 2b and Supplementary Fig. 2a).

**Figure 2.**
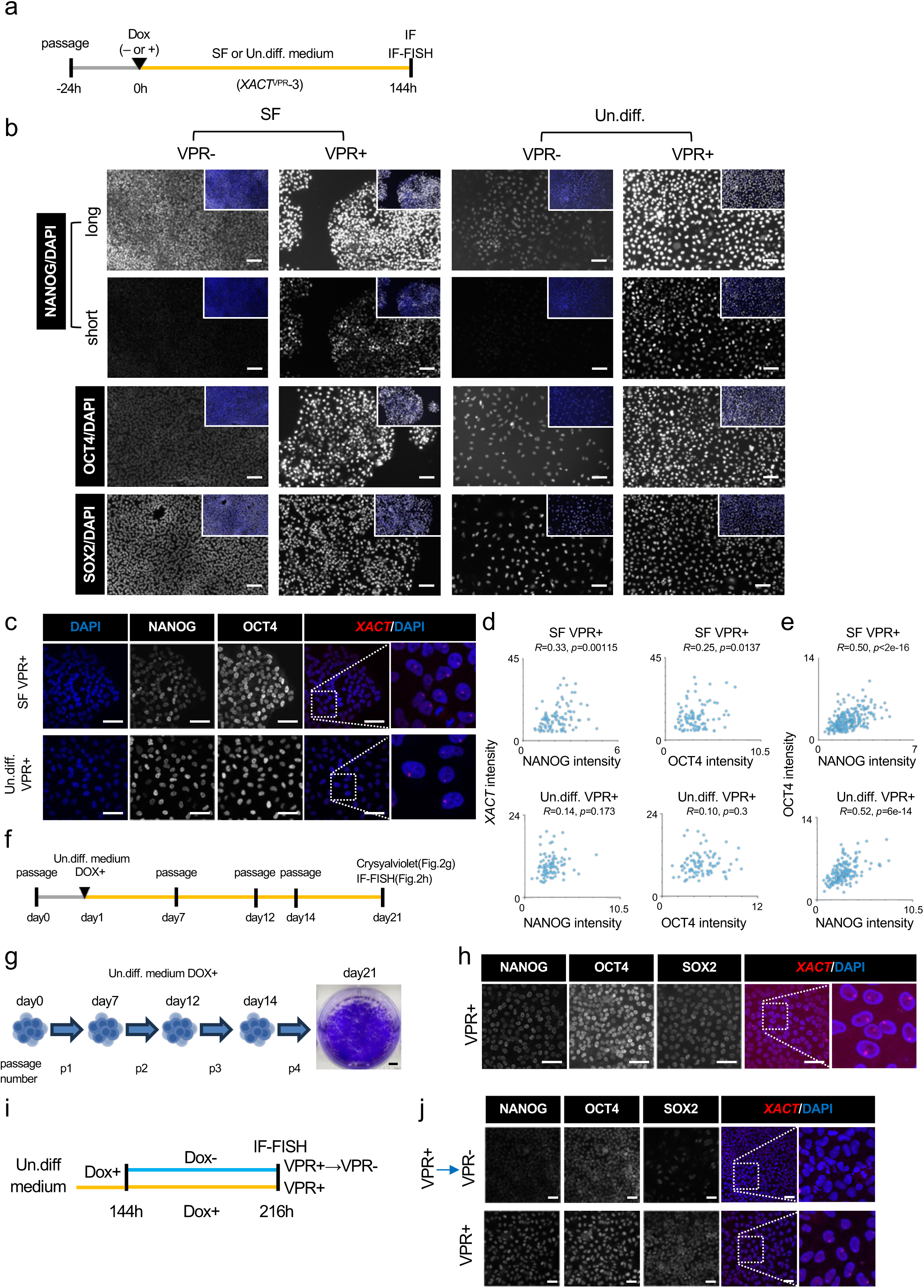
*XACT* upregulation induces heterogeneous and reversible OCT4 and NANOG expression. (a) Experimental scheme for immunofluorescence (IF) and combined IF–RNA-FISH (IF-FISH) using *XACT*^VPR-3^ cells cultured under SF or Un.diff. conditions. Data for *XACT*^VPR^-6 are shown in Supplementary Fig. S2. (b) IF analysis of core pluripotency factors. NANOG was imaged with different exposure times (long and short) to visualize heterogeneity. DAPI was used for nuclear staining. The merge image of entire area was shown in white line. Scale bar, 100 µm. (c) Representative IF-FISH images for NANOG/OCT4/ *XACT* in *XACT*^VPR-3 cells cultured in SF or Un.diff. conditions. Scale bar, 20 μm. (d–e) Correlation of *XACT* expression with NANOG or OCT4 (d), and between NANOG and OCT4 (e), at single-cell resolution. Each dot represents one cell. Correlations were calculated using Pearson’s coefficient; significance was determined by two-tailed t-test. (f) Propagation assay. *XACT*^VPR^-3 cells were maintained in Un.diff. conditions, passaged with TrypLE, and subjected to growth analysis (g) and IF-FISH (h). (g) Crystal violet staining of cell growth. Scale bar, 5mm. (h) IF-FISH of *XACT* and core pluripotency markers. Scale bar, 20 µm. (i) Reversibility assay. *XACT*^VPR^-3 cells were cultured with Dox for 144 h, followed by 72 h of Dox withdrawal, and then analyzed by IF-FISH for *XACT* and core pluripotency factors. Data for *XACT*^VPR^-6 are provided in Supplementary Fig. S2. (j) IF-FISH images (*XACT*/NANOG/OCT4/SOX2) with or without Dox. Scale bar, 50 µm.

Next, we asked whether *XACT* expression itself correlates with the protein hyper elevation of NANOG and OCT4 at the single-cell level. IF combined with RNA-FISH (IF-FISH) revealed that NANOG and OCT4 protein intensities showed only weak correlation with *XACT* RNA abundance (r² = 0.09–0.33 in SF; 0–0.14 in Un.diff.; Fig. 2c,d; Supplementary Fig. S2b,c). By contrast, NANOG and OCT4 proteins were consistently and positively correlated with each other (r² = 0.50–0.58; Fig. 2c,e; Supplementary Fig. S2b,d). These findings indicate that the heterogeneous hyper elevation of NANOG and OCT4 proteins is not directly proportional to *XACT* RNA levels detectable by confocal microscopy. Instead, the coordinated hyper elevation suggests that *XACT* acts through a shared post-transcriptional regulatory mechanism that simultaneously drives both factors.

We then asked whether the pluripotency phenotype induced by *XACT* overexpression could be maintained across multiple passages under Un.diff. conditions, and whether it was reversible. As outlined in the experimental scheme (Fig. 2f), *XACT*^VPR+^ cells were cultured in Un.diff. medium for 20 days with three passages. Proliferation and pluripotency factor expression were assessed by crystal violet staining and IF, respectively. *XACT*^VPR+^ cells proliferated robustly (Fig. 2g and Supplementary Fig. S2e) and maintained heterogeneous expression of core pluripotency factors under these conditions (Fig. 2h and Supplementary Fig. S2f).

To test reversibility, Dox was removed after 144 hours, and cells were cultured for an additional 72 hours in Un.diff. medium without Dox before IF-FISH analysis (Fig. 2i). Upon Dox withdrawal, expression of NANOG, OCT4, SOX2, and *XACT* was markedly reduced (Fig. 2j and Supplementary Fig. S2g).

Together, these results demonstrate that *XACT* upregulation drives heterogeneous hyper elevation of NANOG and OCT4 proteins in both SF and Un.diff. conditions. This phenotype can be stably propagated for extended passages in the absence of bFGF/Activin, yet remains fully reversible once *XACT* is repressed, demonstrating that this phenotype is entirely dependent on sustained *XACT* activity

### *XACT* upregulation impedes repression of pluripotency factors and blocks ectoderm differentiation during directed lineage specification

We next asked whether sustained *XACT* activation could prevent lineage specification under small-molecule–mediated directed differentiation. *XACT*^VPR^ cells were exposed to induction cocktails for ectoderm, mesoderm, and endoderm lineages (Fig. 3a). Cell growth was first assessed following lineage-specific induction. In contrast to the robust proliferation observed under Un.diff. conditions (Fig. 1e and Supplementary Fig. S1e), crystal violet analysis revealed markedly reduced expansion of *XACT*^VPR+^ cells compared to *XACT*^VPR-^ controls across all three differentiation lineages (Fig. 3b and Supplementary Fig. 3a), suggesting that *XACT* activation compromises cell growth in response to lineage-specific cues.

**Figure 3.**
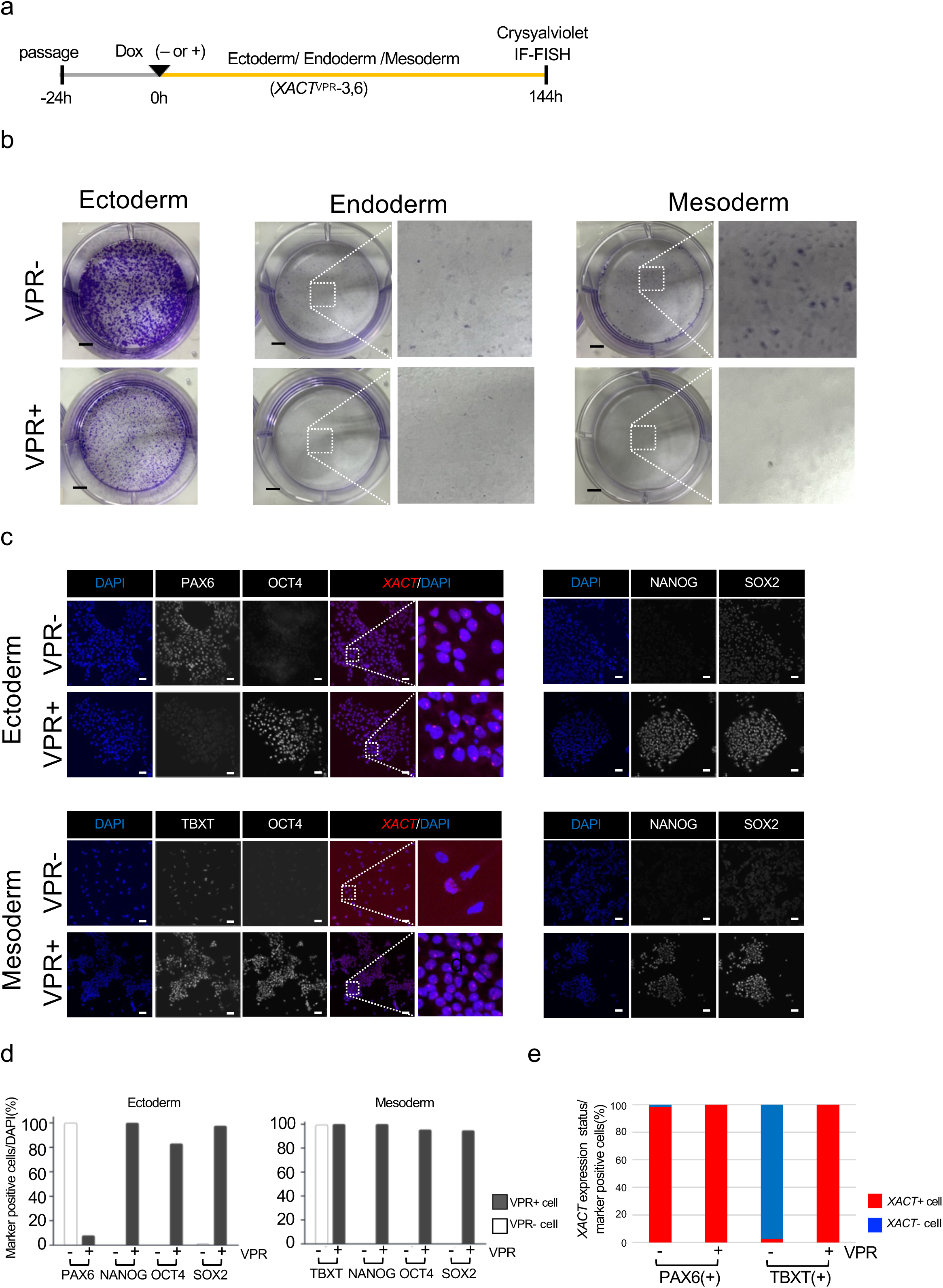
Continuous *XACT* expression impairs directed differentiation of hPSCs. (a) Experimental scheme using *XACT*^VPR^-3 cells for directed differentiation with or without Dox. Differentiation status was assessed by crystal violet staining (b) and IF-FISH (c). Data for *XACT*^VPR^-6 are shown in Supplementary Fig. S3. (b) Crystal violet staining of cells cultured in lineage-specific differentiation media. Scale bar, 5 mm (c) IF-FISH analysis of core pluripotency factors and lineage markers (ectoderm: PAX6; mesoderm: TBXT). Endoderm differentiation was not achieved due to insufficient cell numbers. DAPI was used for nuclear staining. Scale bar, 50 µm. (d) Quantification of marker-positive cells. At least 164 cells were analyzed. (e) Quantification of *XACT* expression in relation to each marker.

We next examined differentiation status in the surviving cells by IF-FISH for lineage markers, core pluripotency factors (NANOG, OCT4, and SOX2), and *XACT* at 144 hours post-induction. Under ectoderm differentiation, nearly all *XACT*^VPR-^ cells expressed PAX6, with 92.1% of these PAX6+ cells also showing detectable *XACT* expression (Fig. 3c,e and Supplementary Fig. S3b,d). In contrast, <10% of *XACT*^VPR+^ cells were PAX6+, whereas >75% retained strong expression of core pluripotency markers (Fig. 3c,d and Supplementary Fig. S3b,c).

In mesoderm differentiation, TBXT was detected in nearly all *XACT*^VPR− cells and in >48.1% of *XACT*^VPR+^ cells (Fig. 3c,d and Supplementary Fig. S3b,c). Notably, in TBXT+ cells, *XACT* RNA was almost entirely absent in *XACT*^VPR-^ cells (97.3%), whereas it remained detectable in the majority of *XACT*^VPR+^ cells (Fig. 3c,e and Supplementary Fig. S3b,c). Endoderm induction was inefficient in both groups, yielding insufficient numbers for meaningful analysis. Together, these findings demonstrate that sustained *XACT* overexpression during directed differentiation impedes exit from pluripotency, most strikingly blocking ectodermal commitment, while allowing partial mesodermal specification. Core pluripotency factors were consistently maintained in both contexts. Considering that additional *XACT* expression under SF conditions drives hyper-elevation of NANOG and OCT4 (Fig. 1h and Supplementary Fig. S1g), and that PAX6 induction can occur irrespective of basal *XACT* expression, we conclude that *XACT* is not strictly required for pluripotency maintenance during directed differentiation. Rather, *XACT* dosage emerges as a key determinant of pluripotency factor levels and lineage commitment capacity.

### *XACT* upregulation generates naïve- and seminoma-like states in a culture context-dependent manner

Because *XACT*^VPR+^ cells proliferate robustly under Un.diff. conditions while maintaining core pluripotency factor expression (Fig. 1e), we next investigated the molecular characteristics associated with altered *XACT* expression. We performed scRNA-seq analysis on *XACT*^WT/Y^, *XACT*^Δ/Y^, *XACT*^VPR-^, and *XACT*^VPR+^ cells under both SF and Un.diff. conditions (Fig. 4a). UMAP analysis revealed that cellular clustering was primarily determined by *XACT* expression status rather than culture condition (Fig. 4b(i–ii)). Cell cycle analysis showed that *XACT*^VPR+^ cells exhibited an increased G1 population compared with *XACT*^WT/Y^, *XACT*^Δ/Y^, and *XACT*^VPR-^ cells in both SF and Un.diff. conditions (Supplementary Fig. 4a,b), consistent with slower proliferation rates (Supplementary Fig. 4b). These data suggest that *XACT* upregulation produces cells distinct from typical hPSCs, which normally display a short G1 phase^22,23^.

**Figure 4.**
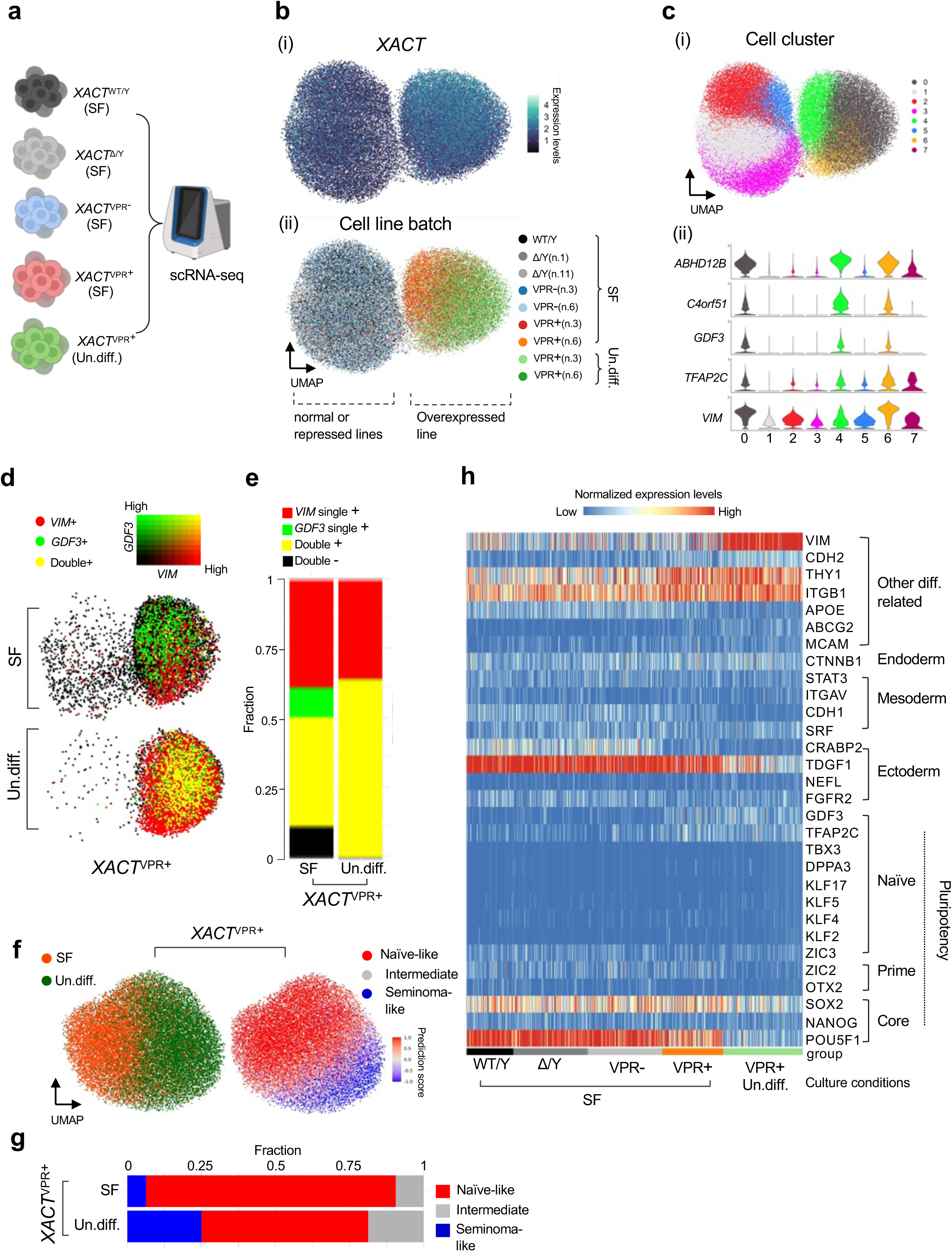
Single-cell transcriptomic profiling of *XACT*-manipulated cells. (a) Experimental design for scRNA-seq using *XACT*^WT/Y^, *XACT*^Δ/Y^, and *XACT*^VPR^ cells. (b) UMAP visualization by *XACT* expression (i) and by genotype/culture condition (ii). (c) UMAP showing cluster identities (i) and marker gene expression (ii): differentiation-defective (ABHD12B, C4orf51), naïve-related (GDF3, TFAP2C), and differentiation (VIM). (d) Co-expression analysis of GDF3 and VIM in *XACT*^VPR+^ cells under Un.diff. conditions. (e) Quantification of co-expressing cells by condition. (f) UMAP of cell-type prediction (naïve-like, seminoma-like, intermediate) in *XACT*^VPR^ cells. Reference cells were derived from integrated datasets of naïve derivatives and seminoma (Supplementary Fig. S5d). (g) Proportions of predicted cell types. (h) Heatmap showing representative pluripotency- and differentiation-associated gene expression.

To identify genes driving these transcriptional states, we extracted differentially expressed genes (DEGs) from clusters 0, 4, and 6 in *XACT*^VPR+^ cells (Fig. 4c(i); Supplementary Fig. 4c; Supplementary Table S1). Notably, naïve pluripotency-associated genes, including GDF3 and TFAP2C^24,25^, were enriched, along with differentiation-defective markers such as ABHD12B and C4orf5^26^ in these clusters (Fig. 4c(ii)). Differentiation-associated genes, including VIM, were also upregulated, particularly in Un.diff. conditions (Fig. 4c(ii); Supplementary Fig. 4d(i)).

Co-expression analysis revealed that a substantial fraction of *XACT*^VPR+^ cells co-expressed GDF3 and VIM, depending on culture conditions: 39% in SF and 64% in Un.diff. (Fig. 4d and 4e). Similarly, DPPA3, a naïve pluripotency marker^27,28^, was predominantly expressed in SF-cultured *XACT*^VPR+^ cells (Supplementary Fig. 4d(ii)), highlighting the context-dependent transcriptional programs induced by *XACT*.

Gene ontology (GO) analysis of DEGs in *XACT*^VPR+^ cells revealed significant downregulation of cellular component (CC) and molecular function (MF) categories in *XACT*^VPR+^ cells (Supplementary Fig. 4e). Notably, junction-related genes (CC) were reduced, potentially explaining the altered morphology observed in *XACT*^VPR+^ cells (Fig. 1d). Within the MF category, histone-binding genes, including CBX2 and PHC1—components of canonical Polycomb repressive complex 1 (PRC1)^29^—were downregulated, suggesting that PRC1 dysfunction may contribute to aberrant activation of differentiation genes such as VIM.

We also observed upregulation of metal ion- and zinc-response genes, including metallothionein (MT) family members, in *XACT*^VPR+^ cells (Supplementary Fig. 4e: GO). Notably, previous studies reported that hESCs exhibiting differentiation failure show elevated MT expression^30^, providing molecular support for the differentiation-defective phenotype observed in *XACT*^VPR+^ cells. Interestingly, high MT expression has also been reported in seminoma and in iPSCs derived from testicular germ cell tumor (TCGT) samples, which frequently exhibit NANOG locus duplication^31^, reflecting a parallel to the hyperactivation of pluripotency factors seen upon *XACT* overexpression. These observations prompted us to investigate whether *XACT* overexpression induces cancer-like transcriptional programs in addition to naïve pluripotency features.

To address the possibility, we checked *XACT* expression status in The Cancer Genome Atlas (TCGA). Surprisingly, *XACT* expression is uniquely and highly expressed in testicular germ cell tumor (Supplementary Fig. 5a), suggesting the similarity of transcriptomic feature of *XACT*^VPR+^ cells to seminoma. To examine this, we analyzed scRNA-seq datasets of and seminoma cells^31^ as well as naïve hPSCs and their derivatives^32^, which showed GDF3 and TFAP2C observed in *XACT*^VPR+^ cells (Fig. 4c (ii)). UMAP analysis revealed clear segregation of these cell types (Supplementary Fig. S5b(i)), with seminoma forming six distinct clusters (clusters: 0,3,6,10,11, and 13 Supplementary Fig. S5b(ii)) and naïve cells clustering according to culture conditions (Supplementary Fig. S5b(i)). Notably, *XACT* expression was enriched in cluster 6 of seminoma, which also exhibited high NANOG and GDF3 levels (Supplementary Fig. S5b(i) and Fig. S5c). In naïve cells and their derivatives, NANOG and GDF3 were robustly expressed in clusters 2, 3, and 5 (Supplementary Fig. S5b(i) and Fig. S5c). Using these clusters as references, we assessed the similarity of *XACT*^VPR+^ cells by prediction scoring (Supplementary Fig. S5d). In SF conditions, 84% of *XACT*^VPR+ cells were classified as naïve-like, whereas this population decreased to 56% under Un.diff. condition. Conversely, seminoma-like cells comprised 6% in SF and 25% in Un.diff. conditions, respectively (Fig. 4f and 4g). These results indicate that the transcriptomic state of *XACT*^VPR+^ cells is context-dependent, and that a subset acquires cancer-like features, highlighting a potential novel role for *XACT* in tumor biology.

We further evaluated the expression of additional pluripotency- and differentiation-associated genes. Heatmap analysis showed that other naïve markers, such as KLF2 and ZIC3^28^, as well as lineage-specific genes including FGFR2 (ectoderm)^33,34^, CDH1 (mesoderm)^33,35^, and CTNB1 (endoderm)^33,36^, were not significantly upregulated in *XACT*^VPR+^ cells under either SF or Un.diff. conditions (Fig. 4h). This indicates that *XACT* overexpression does not drive activation of specific lineage programs.

Together, these findings demonstrate that *XACT* overexpression generates a unique cell population with both naïve pluripotency- and seminoma-like transcriptional features in a culture context–dependent manner.

### *XACT* upregulation causes repression of 3**′** UTR transcripts of OCT4 and NANOG

One unresolved question in this study is the mechanism underlying the differential mRNA and protein expression of core pluripotency factors (Fig. 1g and 1h). Surprisingly, scRNA-seq analysis revealed that OCT4 transcript levels were reduced in *XACT*^VPR+^ cells, particularly under Un.diff. conditions (Fig. 4h). Similarly, NANOG transcripts were not upregulated in *XACT*^VPR+^ cells under Un.diff., inconsistent with qPCR and western blot data (Fig. 1g and 1h). As scRNA-seq predominantly captures 3’ transcript ends, including 3’ untranslated regions (UTRs)^37^, and given that UTRs can influence protein expression^38^, we hypothesized that *XACT* upregulation might selectively alter 3’ transcriptional status, resulting in the observed discrepancies between mRNA and protein levels.

To test this possibility, we analyzed read mapping in scRNA-seq data. Intriguingly, for OCT4, reads corresponding to the 3’ UTR were moderately and markedly reduced in *XACT*^VPR+^ cells under SF and Un.diff. conditions, respectively (Fig. 5a). NANOG transcripts at the 3’ UTR were strongly repressed in *XACT*^VPR+^ cells under both conditions, whereas coding sequence (CDS) reads were enriched (Fig. 5a). In contrast, SOX2 transcript mapping was largely unchanged between *XACT*^VPR+^ and *XACT*^VPR-^ cells (Fig. 5a).

**Figure 5.**
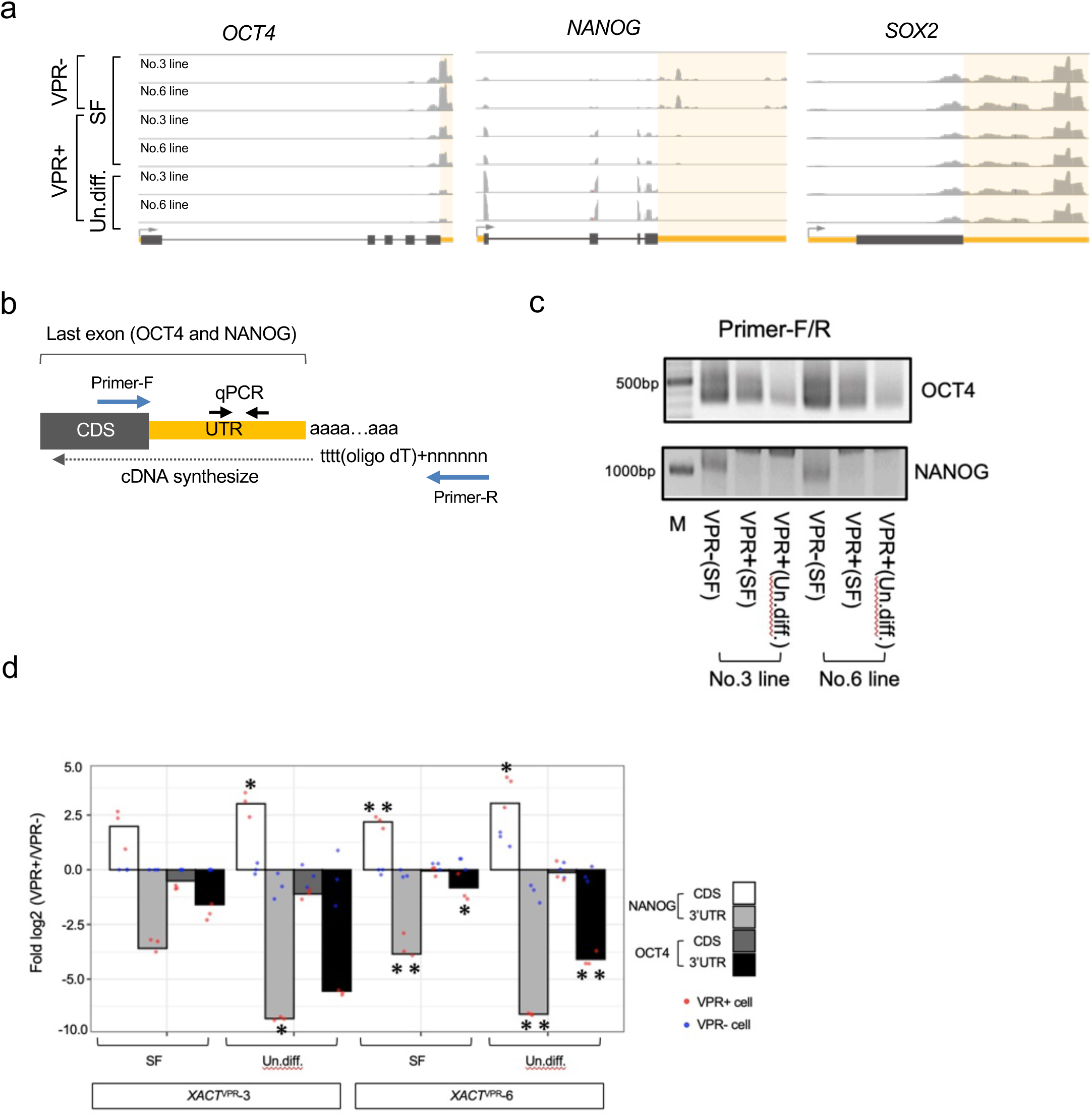
***XACT* upregulation represses 3**′ **UTRs of NANOG and OCT4.** (a) Read coverage of pluripotency factor transcripts in *XACT*^VPR^ cells (scRNA-seq). UTR regions are highlighted in yellow. (b) Experimental design for 3’ RACE. Primer positions are indicated; qPCR primers targeting exon junctions within coding sequence (CDS) and 3’ UTR were used. (c) RT-PCR results for OCT4 and NANOG (full gels in Supplementary Fig. S6a). (d) qPCR analysis of OCT4 using cDNA synthesized with oligo-dT primers. Bar graph shows mean fold-change values (log□) from three biological replicates, with individual data points overlaid. *P* < 0.05; \**P* < 0.01 (Student’s *t*-test).

We next validated 3’ UTR repression using 3’ rapid amplification of cDNA ends (3’ RACE) (Fig. 5b). RT-PCR showed reduced amplification of the OCT4 3’ UTR in *XACT*^VPR+^ cells, particularly under Un.diff. conditions (Fig. 5c). NANOG 3’ UTR products were undetectable in *XACT*^VPR+^ cells under both SF and Un.diff. conditions (Fig. 5c). Unexpectedly, we observed long RT-PCR products for both genes, but Sanger sequencing revealed these to be non-specific (Supplementary Fig. S6a). Genomic PCR confirmed that there were no deletions in the 3’ UTR regions of OCT4 or NANOG (Supplementary Fig. S6b), excluding genomic loss, alternative splicing, or alternative polyadenylation as causes for 3’ UTR repression.

For robust quantification, we performed TaqMan assays targeting 3’ UTR sequences using oligo-dT–primed cDNA (Fig. 5d). This analysis revealed that the 3’ UTR transcripts of NANOG and OCT4 were markedly reduced in *XACT*^VPR+^ cells compared to *XACT*^VPR-^ cells, whereas CDS-derived transcripts showed a distinct pattern. For NANOG, 3’ UTR levels were reduced by approximately 6–10-fold under SF conditions and by 50–70-fold under Un.diff. conditions (Fig. 5d), while CDS levels were instead elevated by ∼3.6–4.8-fold under SF and by ∼5–8-fold under Un.diff. conditions (Fig. 5d). For OCT4, 3’ UTR levels were reduced by ∼2-fold under SF and by 15–47-fold under Un.diff. conditions, yet CDS levels remained largely unchanged (Fig. 5d, 0.55–0.93-fold across both conditions).

Together, these results demonstrate that *XACT* upregulation selectively represses 3’ UTR transcripts of OCT4 and NANOG in both SF and Un.diff. conditions, providing a molecular basis for the observed protein hyper elevation and heterogeneous expression of these core pluripotency factors.

### *XACT*-high female hPSCs exhibit transcriptional features resembling *XACT*^VPR+ cells

Previous studies have reported that *XACT* overexpression is a hallmark of X-chromosome inactivation (XCI) erosion, typically caused by the loss of XIST expression in female hPSCs^15,16,19,39^. We therefore asked whether female hPSCs with eroded XCI display transcriptional features similar to *XACT*^VPR+^ cells in SF conditions. Reanalysis of bulk RNA-seq data from partially-eroded and eroded female iPSCs (Supplementary Fig. S7a) revealed elevated expression of NANOG, GDF3, DPPA3, and TFAP2C in eroded cells, closely resembling the transcriptomic profile of *XACT*^VPR+^ cells (Supplementary Fig. S7b).

To further characterize this relationship, we reanalyzed public RNA-seq data from 106 female hPSC lines in the HipSci project^40^. We observed a significant negative correlation between *XACT* and XIST expression (Pearson R = –0.286, p = 0.0029; Supplementary Fig. S7c). Based on expression levels of these two genes, 49 lines were selected for further analysis and classified as either *XACT*-high (XIST-low) or *XACT*-low (XIST-high) (Supplementary Fig. S7c). Strikingly, *XACT*-high lines exhibited significantly elevated expression of pluripotency- and naïve-associated genes, including NANOG, OCT4, GDF3, DPPA3, and TFAP2C, as well as differentiation-related markers such as HAND1 and GATA6 (Supplementary Fig. S7d).

Together, these findings demonstrate that *XACT*-high female hPSCs, including XCI-eroded lines, share transcriptional features with *XACT*^VPR+^ cells. This resemblance provides a potential molecular explanation for the impaired developmental potential of XCI-eroded female hPSCs^19^, implicating aberrant upregulation of pluripotency- and naïve-related genes as a contributing factor.

### No evidence of genomic proximity between GDF3/DPPA3/NANOG and the relaxed *XACT* locus

A notable feature of *XACT*^VPR+^ cells was the concomitant upregulation of GDF3, DPPA3, and NANOG under SF conditions (Fig. 4h; Supplementary Fig. 4d (ii)). Since these genes are clustered within the chr12p13.31 region (Supplementary Fig. 8a), we hypothesized that *XACT* activation might alter higher-order chromatin conformation and thereby influence their expression.

To test this, we performed three-dimensional DNA-FISH targeting the *XACT* locus and the GDF3/DPPA3/NANOG cluster, followed by super-resolution microscopy in pluripotent conditions. Consistent with dCas9-VPR–mediated transcriptional activation, we observed significant chromatin relaxation at the *XACT* locus in *XACT*^VPR+^ cells compared with *XACT*^VPR-^ controls (Supplementary Fig. 8b). However, no relaxation was detected at the GDF3/DPPA3/NANOG loci, nor did we observe any increased spatial proximity between these loci and *XACT* (Supplementary Fig. 8b).

These results demonstrate that the co-upregulation of GDF3, DPPA3, and NANOG upon *XACT* upregulation is not explained by direct chromatin decompaction or physical proximity to the *XACT* locus, suggesting that their induction is instead mediated through downstream regulatory networks or indirect mechanisms.

### Prolonged *XACT* expression impairs post-implantation modeling

Our findings that *XACT* upregulation disrupts both undirected and directed differentiation prompted us to ask whether its persistence also interferes with early human embryogenesis. Because *XACT* is uniquely expressed in pre-implantation embryos and normally repressed after implantation^3,15^, we reasoned that inappropriate maintenance of *XACT* expression could impair post-implantation developmental programs.

To address this, we first generated post-implantation embryo models—axioloids^41^ and embryoid bodies (EBs)—and monitored *XACT* dynamics during their formation (Fig. 6a). RNA-FISH revealed that *XACT* expression persisted for up to 48 hours in EBs and in 2D cultures treated with CHIR99021 and bFGF, conditions used to promote axioloid formation (Fig. 6b). In contrast, developing axioloids showed marked repression of *XACT*, consistent with previous reports of its downregulation around implantation in human embryos^3,15^. These data confirm that in vitro–generated axioloids recapitulate the silencing of *XACT* during the implantation transition.

**Figure 6.**
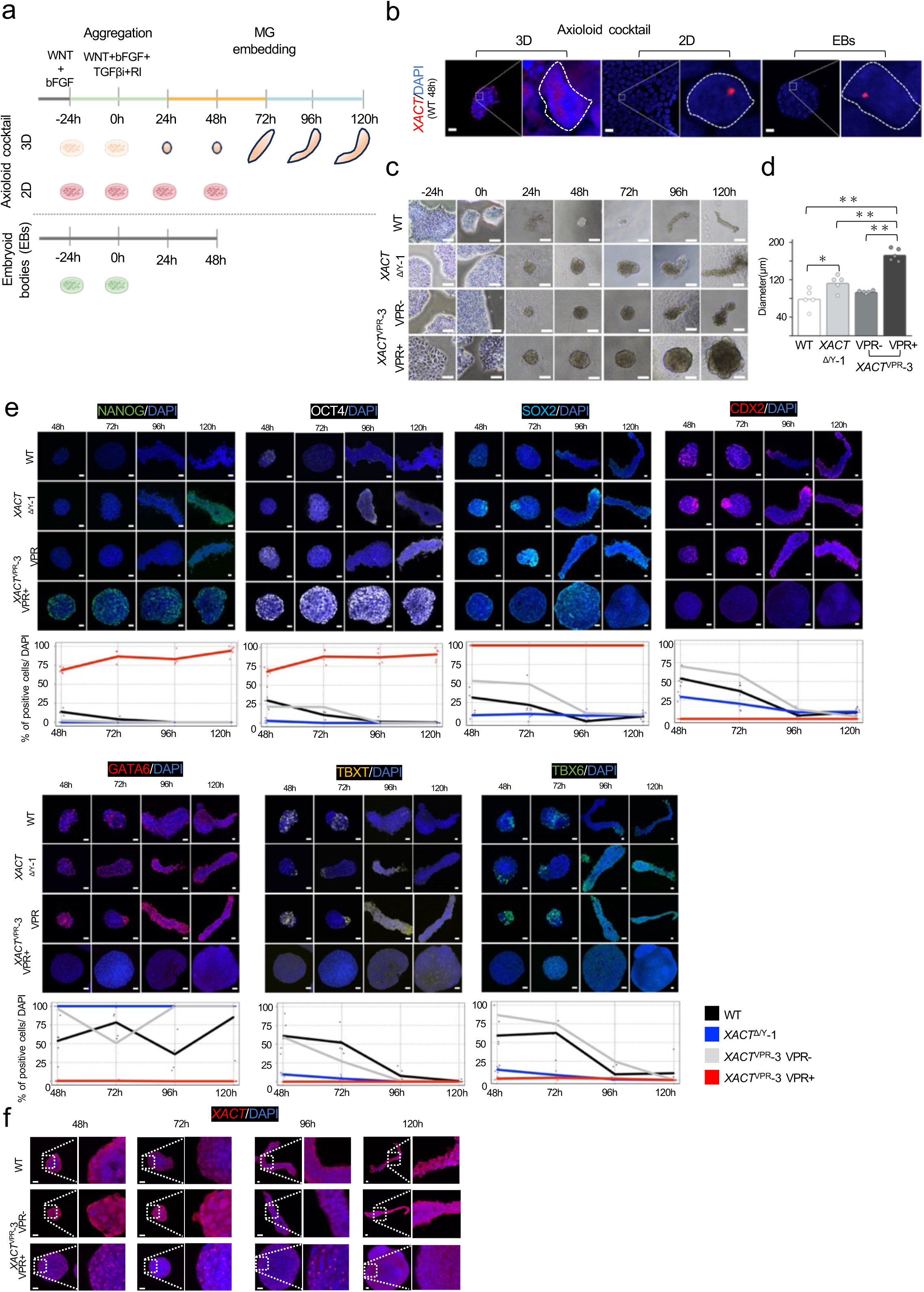
Prolonged *XACT* expression disrupts post-implantation modeling. (a) Experimental scheme profiling *XACT* expression during post-implantation modeling using three approaches: 3D Axioloid cocktail, 2D Axioloid cocktail, and embryoid body (EB) formation. (b) RNA-FISH of *XACT* expression at 48 h post-aggregation. DAPI-stained nuclei are outlined in white. Scale bar, 20 µm. (c) Bright-field images of axioloid morphology. Scale bar, 100 µm. (d) Quantification of axioloid diameter across genotypes at 48 h. *P < 0.05; **P < 0.01. Statistical significance was determined by one-way ANOVA with Tukey’s multiple-comparison test. A significant overall effect was followed by post hoc pairwise comparisons with appropriate multiple-testing correction. (e–f) IF-FISH analysis of *XACT* and lineage markers in axioloids. (e) IF images for core pluripotency (NANOG, OCT4, SOX2), trophectoderm (CDX2), primitive endoderm (GATA6), and mesoderm (TBXT, TBX6). Percentages of marker-positive cells are plotted. Scale bar, 20 µm. (f) Representative *XACT* RNA-FISH. Scale bar, 20 µm.

We next examined the consequences of sustained *XACT* expression by generating axioloids from WT, *XACT*^Δ/Y^, and *XACT*^VPR^ lines. By 48 hours post-aggregation, *XACT*^VPR+^ cells formed significantly larger and more spherical aggregates compared to controls (Fig. 6c–d; Supplementary Fig. S9a and S9b). Whereas WT, *XACT*^Δ/Y^, and *XACT*^VPR-^ cells elongated into asymmetric axioloid-like structures by 96–120 hours, *XACT*^VPR+^ cells failed to undergo this morphological transition and instead retained compact spherical morphologies (Fig. 6c, 6d, Supplementary Fig. S9a, and S9b), indicating that prolonged *XACT* expression disrupts post-implantation morphogenesis.

To further dissect the impact of *XACT* on lineage specification, we performed time-course IF-FISH for pluripotency and differentiation markers. In WT, *XACT*^Δ/Y^, and *XACT*^VPR-^ cells, pluripotency factors (NANOG, OCT4, and SOX2) declined by 96 hours (0–7.7%; Fig. 6e and Supplementary Fig. S9c), coinciding with elongation of the structures. In parallel, lineage markers including CDX2 (trophoblast)^28,33^ and TBXT/TBX6 (mesoderm)^28,33^ emerged as early as 48 hours (CDX2: 25.2–54.1%; TBXT: 9.8-61.0%; TBX6: 13.9-63.4%) and showed asymmetric expression by 120 hours (Fig. 6e and Supplementary Fig. S9c). GATA6 (primitive endoderm)^28,33^ was also robustly induced, with expression observed throughout axioloid-like structures by 120 hours (85.5–100%; Fig. 6e and Supplementary Fig. S9c).

In contrast, *XACT*^VPR+^ cells retained high NANOG and OCT4 expression across the majority of cells (90.7–100%; Fig. 6e,f and Supplementary Fig. S9c,d). SOX2 expression, however, was variable after 96 hours, and lineage markers failed to be induced under these conditions. Notably, *XACT* expression itself was consistently maintained in most cells (Fig. 6e,f and Supplementary Fig. S9c,d).

Together, these findings demonstrate that inappropriate maintenance of *XACT* expression impedes post-implantation modeling by locking cells in a pluripotent state. This suggests that dysregulation of *XACT*, a normally pre-implantation–restricted lncRNA, may represent a mechanism underlying developmental failure in early human embryos.

### *XACT* expression positively correlates with pluripotent cells in human pre- and post-implantation embryos

Finally, to evaluate whether *XACT* expression in vivo is associated with pluripotency networks, we reanalyzed published single-cell datasets from human embryos spanning pre- to post-implantation stages^28,42^. We focused on the relationship between *XACT* and previously defined pluripotency- and differentiation-related markers. Across both pre- and peri-implantation datasets, the three genes most positively correlated with *XACT* were GDF3, POU5F1(OCT4), and NANOG, whereas differentiation markers such as GATA3 and KRT7 showed negative correlations (Fig. 7a; Supplementary Table S2).

**Figure 7.**
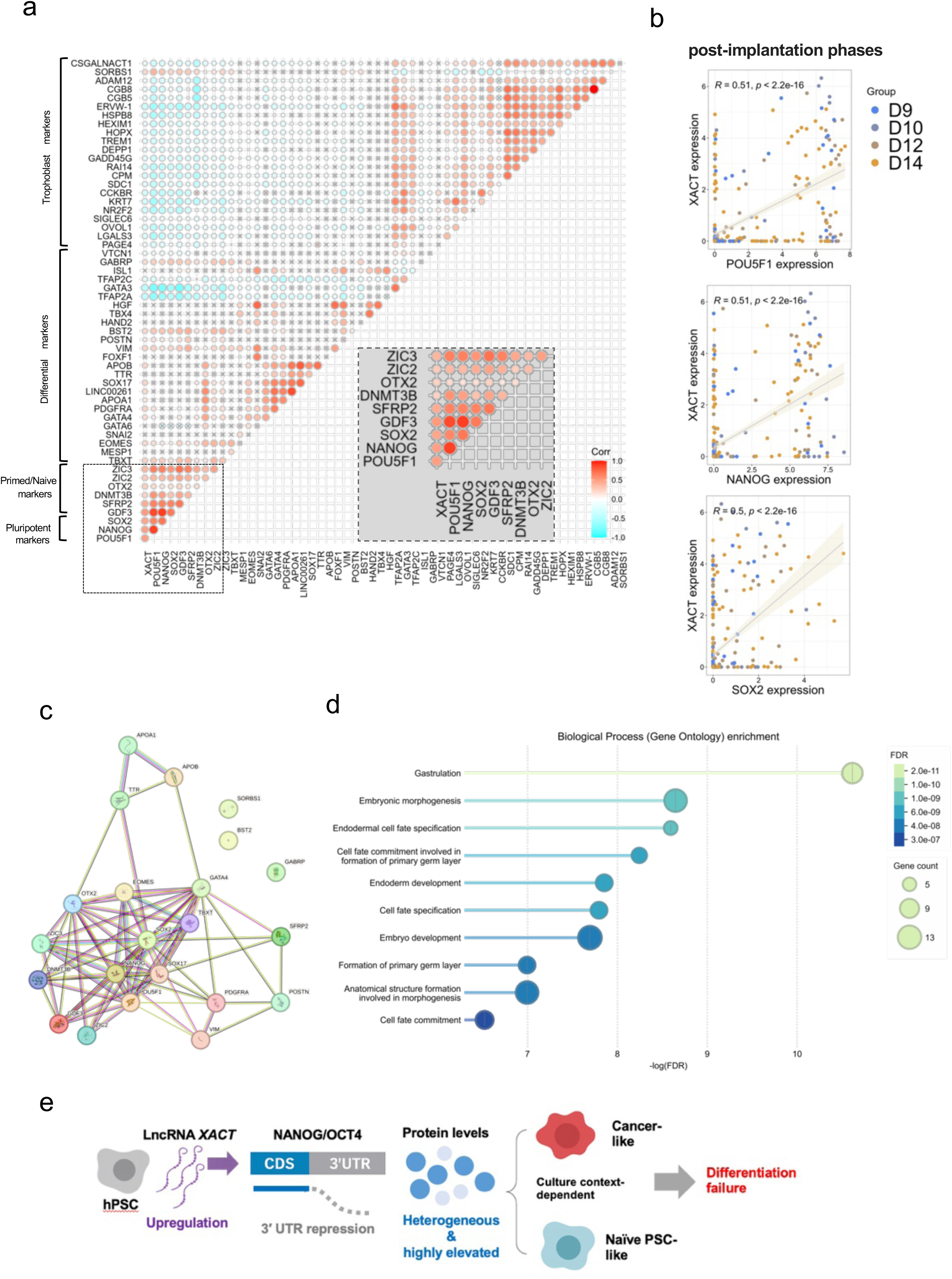
*XACT* correlates with the pluripotency network in human embryos. (a) Correlation matrix of pluripotency and differentiation markers with *XACT* in pre- and post-implantation embryos (GSE136477). Correlations between *XACT* and pluripotency-related genes are highlighted. Significant correlations (P < 0.01) are shown as colored circles. (b) Correlation of *XACT* with core pluripotency gene expression in post-implantation embryonic cells (Day 9–14). Each dot represents a single cell. (c) Protein–protein interaction (PPI) network of genes positively correlated with *XACT*, generated by STRING. Core pluripotency factors formed a central cluster (PPI enrichment p < 1.0e-16). (d) Gene Ontology (GO) enrichment analysis of the PPI gene set. Top 10 biological processes with FDR values are shown. (e) Model of *XACT* function in human embryogenesis. *XACT* is normally expressed in pluripotent cells and downregulated upon differentiation. Loss of *XACT* permits normal pluripotency maintenance and differentiation. In contrast, persistent or ectopic *XACT* expression drives simultaneous activation of naïve-like pluripotency and differentiation genes, leading to culture context–dependent outcomes: naïve-like transcriptome under bFGF/Activin conditions, or tumor-like transcriptome without these factors, thereby impairing lineage commitment and developmental progression.

To determine whether this association persists beyond implantation, we analyzed datasets from day 9–14 embryos, corresponding to early post-implantation stages. Strikingly, *XACT* expression continued to show positive correlations with core pluripotency factors, including POU5F1(OCT4), NANOG, and SOX2 (Fig. 7b). We next asked whether *XACT*-correlated genes during post-implantation stages constitute a functional network. STRING analysis^43^ revealed a highly significant protein–protein interaction (PPI) network centered on core pluripotency factors (PPI enrichment p < 1.0e-16; Fig. 7c), and GO analysis demonstrated enrichment for biological processes related to embryonic development (Fig. 7d; Supplementary Table S2).

Together, these results show that *XACT* expression remains tightly linked to pluripotency-associated transcriptional programs both before and after implantation. This finding suggests that precise spatiotemporal regulation of *XACT* is critical for appropriate early human embryogenesis.

## Discussion

In this study, we identified the lncRNA *XACT* as a critical modulator of cell fate decisions in hPSCs. Our findings highlight that misregulation of a developmentally restricted lncRNA can profoundly alter pluripotent cell identity in a culture context-dependent manner—promoting naïve-like states in standard hPSC conditions, driving tumor-like programs in the absence of exogenous factors, and blocking lineage specification (Fig. 7e). This places *XACT* among a very limited subset of lncRNAs capable of reshaping genome-wide transcriptional programs.

*XACT* was originally described as an active X-chromosome transcript with expression restricted to hPSCs, displaying a reciprocal pattern to the XCI master regulator XIST^17^. Unlike XIST, which mainly acts in cis, our findings suggest that *XACT* functions primarily in trans, similar to other regulatory lncRNAs such as HOTAIR^44^, TUNA^45^, Linc-ROR^46^, and lincRNA-p21^47^. Among these, TUNA and Linc-ROR regulate pluripotency through distinct mechanisms: TUNA interacts with the hnRNP-K/NCL/PTBP1 complex to modulate promoters of pluripotency factors in mice^45^, while Linc-ROR acts as a miRNA sponge for miR-145, thereby protecting pluripotency transcripts in hPSCs^46^. Although Linc-ROR upregulation promotes pluripotency factor expression during differentiation, the response appears relatively homogeneous^46^. In contrast, *XACT* activation drives hyperactivation of OCT4 and NANOG proteins accompanied by marked heterogeneity and repression of their 3’UTRs (Fig. 2, and Fig. 5). Given that miRNA-mediated RNA degradation is a widespread mechanism^48^, these findings imply that the mode of action of *XACT* differs fundamentally from that of Linc-ROR.

The precise mechanism underlying *XACT*-mediated OCT4 and NANOG hyperactivation remains to be clarified. Recent evidence suggests that deletion of 3’ UTRs can upregulate CDS expression in human cells (doi: 10.1101/2022.03.15.484504). Considering that 3’ UTR status affects both translational efficiency and mRNA stability^38^, our findings raise the possibility that *XACT*-mediated repression of 3’ UTRs contributes to excessive OCT4 and NANOG protein levels.

At the cellular phenotype level, *XACT* upregulation markedly alters the dependency of hPSCs on extrinsic signals. Typically, maintenance of hPSCs requires supplementation with bFGF/Activin in the culture medium^20^. Remarkably, *XACT* activation compensates for these essential molecules and sustains high levels of pluripotency factors even under minimal conditions (Fig. 2). This suggests that hyper elevation of core pluripotency factors enables hPSCs to overcome their reliance on external signaling.

From another perspective, *XACT* upregulation also expands populations with seminoma-like transcriptional features, raising safety concerns for transplantation medicine if *XACT* were aberrantly activated in environments such as Un.diff. medium. This finding underscores the dual nature of pluripotency regulation: while *XACT* sustains self-renewal, its misregulation can tip cells toward tumor-like states. Importantly, both our present results and previous studies demonstrate that *XACT* deletion does not induce oncogenic programs and, instead, facilitates efficient neuronal differentiation^16^. Thus, targeted removal of *XACT* activity may not only mitigate tumorigenic risk but also improve lineage commitment, providing a promising strategy to enhance the safety of hPSC-based regenerative applications.

Finally, considering the inherent limitations of reverse genetic approaches in human embryos, our study introduces a bi-directional strategy combining hPSCs and embryo modeling to interrogate gene function. This framework provides a powerful tool for uncovering regulators essential for human development. In the future, combining CRISPR-based screening with this bi-directional approach may greatly advance our understanding of molecular networks in early embryogenesis and help identify genetic factors contributing to unexplained miscarriages.

## Methods

### hPSC culture

Male BJ-iPSCs were used for *XACT* functional assays, and the *XACT* knockout (KO) line was previously established ^16^. hPSCs were cultured as previously described ^16,19,49^ with slight modifications. Briefly, hPSCs were maintained in StemFlex medium (Thermo Fisher Scientific, Waltham, MA, USA) supplemented with 50 U/ml penicillin and 50 μg/ml streptomycin (Thermo Fisher Scientific, Waltham, MA, USA) on Matrigel-coated plates (Corning Inc., NY, USA). For routine passaging, hPSCs were dissociated using 1 mM EDTA and maintained in culture medium supplemented with 10 μM Y-27632 (RI: Rho-kinase inhibitor; StemCell Technologies, Vancouver, Canada) for 24 hours. Mycoplasma contamination was routinely tested and confirmed to be negative. For axioloid derivation, hPSCs were cultured in StemFit AK02N medium (ReproCELL, Yokohama, Kanagawa, Japan) on iMatrix-511 silk (Nippi, Kusatsu, Shiga, Japan) for at least 14 days before differentiation.

### Crystal violet staining

Cells were seeded in 6-well plates at a density of 1.25 × 10□ cells per well and cultured for 24 h in StemFlex medium (Thermo Fisher Scientific, Waltham, MA, USA) supplemented with 10 μM Y-27632 (RI). On the following day, the medium was replaced with undirected differentiation medium, directed differentiation medium, or StemFlex medium, depending on the assay. Cells were then maintained for the indicated period in the presence or absence of doxycycline (Dox). For crystal violet staining, cells were fixed with 4% formaldehyde for 25 min at room temperature, washed with PBS (–), and stained with 0.01% crystal violet (MilliporeSigma, Darmstadt, Germany) for 60 min. Excess dye was removed with distilled water, and the stained cells were imaged and quantified using Fiji (ImageJ, NIH).

### Generation of *XACT*^Δ/Y^ and *XACT*^VPR^ lines

The *XACT*^Δ/Y^-1 and *XACT*^Δ/Y^-11 cell lines, derived from BJ-iPSCs, were previously generated and used in this study ^16^. The same parental BJ-iPSCs were also used to generate the *XACT*^VPR+^ cell line. To induce *XACT* expression by doxycycline (Dox), the PB-TRE-dCas9-VPR plasmid (a gift from George Church, Addgene plasmid #63800) and a gRNA-expressing vector within the PiggyBac system, previously utilized for *XACT* knockdown experiments ^16^, were co-transfected along with PBase^16,18^. For transfection, 1 μg of each gRNA (gRNA1 and gRNA2) and 10μg of dCas-TRE-VPR were used. Following transfection, hygromycin B (50 μg/mL, Thermo Fisher Scientific, Waltham, MA, USA) was added to the culture medium, and after one week, puromycin (1 μg/mL, Thermo Fisher Scientific, Waltham, MA, USA) was also introduced. Both antibiotics were continuously maintained in the culture medium throughout the culture period. After subcloning, *XACT* expression upon Dox induction was assessed using RNA-FISH, and two independent clones, *XACT*^VPR^-3 and *XACT*^VPR^-6, were selected for further analysis. The gRNA sequences used for *XACT* activation are listed in Table S3.

### Undirected differentiation(Un.diff.)

Undirected differentiation experiments were performed for 5–6 days on Matrigel- or gelatin-coated dishes by transitioning hPSCs into differentiation medium composed of DMEM/F-12 (Thermo Fisher Scientific, Waltham, MA, USA) supplemented with 20% KnockOut Serum Replacement (KSR, Thermo Fisher Scientific, Waltham, MA, USA), 2 mM GlutaMAX (Thermo Fisher Scientific, Waltham, MA, USA), 0.1 mM non-essential amino acids (NEAA; Thermo Fisher Scientific, Waltham, MA, USA), 10 μM Y-27632 (RI) (STEMCELL Technologies, Vancouver, Canada), and 50 U/ml penicillin and 50 μg/ml streptomycin (Thermo Fisher Scientific, Waltham, MA, USA).

### Directed differentiation

Directed differentiation was performed using the STEMdiff™ Trilineage Differentiation Kit (STEMCELL Technologies, Vancouver, Canada) according to the manufacturer’s instructions. Differentiated cells were subsequently analyzed by immunofluorescence staining and crystal violet staining. The antibodies used for immunofluorescence are listed in Supplementary Table S4.

### Immunofluorescence (IF)

IF staining was conducted following established protocols with slight modifications ^49^. Briefly, cells were fixed in 4% paraformaldehyde (PFA) prepared in PBS for 25 minutes at room temperature, then permeabilized using 0.1% Triton X-100 for another 20 minutes. After washing, nonspecific binding was blocked by incubation with 1.5% BSA in PBS (-) for 1 hour. Cells were incubated sequentially with primary and fluorophore-conjugated secondary antibodies, and imaging was conducted using a Carl Zeiss LSM880 confocal microscope (Zeiss, Jena, Germany) or IX83(OLYMPUS, Japan).

### Immunofluorescence combined with RNA-FISH (immune-FISH)

Immuno-FISH was performed as previously described ^50^. Briefly, cells were cultured on coverslips and fixed in 4% paraformaldehyde in PBS (-) (NACALAI TESQUE, INC., Kyoto, Japan) for 25 minutes at room temperature. The cells were then permeabilized with 0.25% Triton X-100 for 20 minutes and subjected to immunostaining. For blocking, RNaseOUT (Thermo Fisher Scientific, Waltham, MA, USA) was added to 1.5% BSA (VWR International, LLC., Randor, Pennsylvania, USA)-PBS(-) and incubated for 1 hour at room temperature. Primary antibodies were diluted in BSA-PBS containing RNaseOUT and incubated for 1 hour at room temperature. The primary antibodies used in this study included rabbit anti-NANOG (1:100, (Abcam, Tokyo, Japan)), mouse anti-OCT4 (1:300, (Santa Cruz, Dallas, Texas, USA)), goat anti-SOX2 (1:100, (R&D Systems, Inc., Minneapolis, Minnesota, USA)), anti-PAX6(1:300, (Invitrogen™, Tokyo, Japan)), and anti-TBXT(1:100, (Abcam, Tokyo, Japan)). After primary antibody incubation, samples were washed and incubated with secondary antibodies diluted 1:500 in PBS (-). The secondary antibodies used in this study were as follows: Goat anti-rabbit Alexa Fluor 488 (Thermo Fisher Scientific, Waltham, MA, USA), Goat anti-mouse Alexa Fluor 633 (Thermo Fisher Scientific, Waltham, MA, USA), Donkey anti-goat Alexa Fluor 633(Thermo Fisher Scientific, Waltham, MA, USA), and Donkey anti-goat Alexa Fluor 488 (Thermo Fisher Scientific, Waltham, MA, USA). Following immunostaining, RNA-FISH was performed using probes prepared with a nick translation kit (Abbott Laboratories, Chicago, IL, USA). The following probes were used for target detection: XIST: entire-hXIST ^19^, *XACT*: CTD-3063K22 (BAC clone), and HUWE1: PR11-975N19 (BAC clone). Nuclei were counterstained with VECTASHIELD mounting medium containing DAPI (Vector Laboratories, Inc., Newark California, USA). Imaging was conducted using a Carl Zeiss LSM880 confocal microscope (Zeiss, Jena, Germany)or FV3000(OLYMPUS, Japan).

For signal quantification, we used QuPATH (https://qupath.github.io/). Cell boundaries were defined based on DAPI staining, with thresholding parameters optimized for accurate segmentation. Following segmentation, the mean fluorescence intensity of the target channel was measured for each cell using the *Mean Intensity* parameter. This value was then normalized to the DAPI-stained nuclear area, yielding a per-cell intensity measure independent of nuclear size.

### RNA-FISH

Cells were cultured on coverslips and fixed in 4% paraformaldehyde in PBS (-) (NACALAI TESQUE, INC., Kyoto, Japan) for 25 minutes at room temperature. The cells were then permeabilized with 0.25% Triton X-100 for 20 minutes. After that, RNA-FISH was performed using probes prepared with a nick translation kit (Abbott Laboratories, Chicago, IL, USA). The following probes were used for target detection: *XACT*: CTD-3063K22 (BAC clone). Nuclei were counterstained with VECTASHIELD mounting medium containing DAPI (Vector Laboratories, Inc., Newark California, USA). Imaging was conducted using a Carl Zeiss LSM880 confocal microscope (Zeiss, Jena, Germany) or FV3000(OLYMPUS, Japan).

### DNA-FISH

DNA-FISH was performed as previously described ^19,50^. Briefly, fixed and permeabilized cells were treated with RNase A and incubated in 0.2N HCl containing 0.05% Tween-20 on ice for 10 minutes. The samples were then incubated at 85°C for 10 minutes, followed by overnight hybridization at 37°C. Probes for XIST (entire-hXIST), *XACT* (CTD-3063K22), and the NANOG/GDF3/DPPA3 locus RP11-103J24 (BAC clone) were prepared using a nick translation kit. Super-resolution imaging was performed using Airyscan mode on a Carl Zeiss LSM880 microscope equipped with a 64× Plan-Apochromat objective lens (Zeiss, Jena, Germany). For signal distance analysis, the inter-signal distance between two DNA-FISH signals was manually measured using FIJI (ImageJ, NIH). The straight line tool was applied to draw a line from the edge of one fluorescent signal to the edge of the other, and the resulting distance was recorded.

### Western Blotting

Cells were lysed in RIPA buffer supplemented with protease inhibitors, and protein concentrations were determined using a DC protein assay. Equal amounts of protein (5 µg per sample) were subjected to SDS–PAGE and transferred onto PVDF nitrocellulose membranes. Membranes were blocked with 1.5% skim milk and probed with primary antibodies against NANOG, OCT4, SOX2, and GAPDH. After incubation with secondary antibodies, signals were detected using Western Lighting Ultra.

### Quantitative PCR (qPCR)

Total RNA was extracted using the RNeasy Plus Micro Kit (QIAGEN, Hilden, Germany) according to the manufacturer’s instructions. cDNA synthesis was performed using the iScript cDNA Synthesis Kit (Bio-Rad, Hercules, CA, USA) with a mixture of random hexamers and oligo(dT) primers. For oligo(dT)-primed reverse transcription, an anchored oligo(dT) primer (Merck, Darmstadt, Germany) and SuperScript™ IV Reverse Transcriptase (Thermo Fisher Scientific, Waltham, MA, USA) were used.

Gene expression levels were quantified using the ΔΔCt method, with GAPDH as the internal control. Primer sequences used for qPCR and the TaqMan system are listed in Supplementary Table S4.

### 3′ RACE

Total RNA was extracted as described above, and 60 μg was used for reverse transcription (RT). For the RT reaction, an anchored oligo(dT) primer containing a random sequence (5’-AAGCAGTGGTATCAACGCAGAGTACTTTTTTTTTTTTTTTTTTTTTTTVN-3’) was employed. The random sequence served as the reverse primer for RT-PCR, and the resulting amplicons were analyzed by agarose gel electrophoresis. The forward primers used in this assay are listed in Supplementary Table S4.

### scRNA-seq analysis

Single-cell RNA sequencing (scRNA-seq) was performed using the BD Rhapsody system (BD Biosciences, NJ, USA). Gene expression matrices were generated with cwl-runner using the BD Rhapsody pipeline (rhapsody_pipeline_2.2.cwl). RNA sequences were aligned to the human reference genome GRCh38 and quantified using the GENCODE v42 annotation.

All downstream analyses were conducted in R (https://www.r-project.org/) using Seurat^51^ (v5.0). Cells with >200 detected features and <25% mitochondrial content were retained, and genes expressed in at least ten cells were included. For dimensionality reduction, UMAP was applied to human pluripotent stem cell (hPSC) datasets. Data integration was performed using Seurat’s IntegrateLayers function with Harmony-based integration. Differentially expressed genes (DEGs) in *XACT*^VPR+ cells (clusters 0, 4, and 6) were identified using FindMarkers (Seurat) with a threshold of avg_log2FC ≥ 0.5. Gene ontology (GO) enrichment was conducted with the enrichGO package. For co-expression analysis, the blend option of the FeaturePlot function in Seurat was used to visualize the simultaneous expression of two genes. Heatmaps of naïve, primed, and differentiation-associated genes were generated by extracting the expression matrix from the Seurat object and visualizing in R.

For inspection of read mapping, raw scRNA-seq FASTQ files from the BD Rhapsody system were processed with the BD Rhapsody WTA pipeline to generate BAM files. BAM files were visualized using the Integrative Genomics Viewer (IGV, Broad Institute) to examine read alignments and gene locus coverage.

For prediction score analysis, single-cell RNA-seq datasets from GSE166422 (naïve PSCs and derivatives) and GSE256162 (seminoma dataset, GSM8086745) were integrated using Harmony integration implemented in Seurat (v4.4.0). From the integrated UMAP, clusters 2, 3, 5, and 6—positive for GDF3, NANOG, and *XACT*—were extracted as the reference subset (Supplementary Fig. S5d). The query dataset was generated from *XACT*^VPR+^ cells only. Both reference and query datasets were normalized, variable features were identified, and PCA was applied. Transfer anchors were identified using FindTransferAnchors, and prediction scores were computed with TransferData (Seurat). To evaluate relative similarity, per-cell prediction scores for PXGL_n2b27 (naïve PSCs and derivatives) and seminoma were extracted, and a score difference (score_diff) was calculated as:

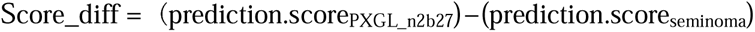

Cells with score_diff ≥ 0.6 were classified as naïve-like (PXGL_n2b27), those with score_diff ≤ −0.6 as seminoma-like, and those between −0.6 and 0.6 as intermediate. UMAP coordinates were extracted from the Seurat object, merged with per-cell score differences, and visualized using ggplot2 (v3.5.1).

For analysis of scRNA-seq data from human pre- and post-implantation embryos, publicly available data from GSE136477 (Xiang et al., 2020) were processed in R. Correlation plots were generated, and Pearson correlation coefficients were calculated to compare gene expression patterns. Genes showing positive correlation with *XACT* in post-implantation cells (Supplementary Table S2) were further analyzed using STRING (https://string-db.org) to explore protein–protein interaction networks.

All raw and processed data generated in this study have been deposited in GEO under accession number GSE300484 (close until the paper is accepted).

### Bulk RNA-seq data analysis

Bulk RNA-seq data of XCI-eroded and partially eroded iPSCs (Supplementary Fig. S7b) were obtained from the Gene Expression Omnibus (GSE218950). Two biological replicates each of eroded and non-eroded hPSCs were analyzed. Gene expression levels were quantified as log□ (TPM + 1) values, and comparisons were performed for a subset of genes associated with naïve pluripotency, primed pluripotency, and differentiation.

For HipSci data set analysis (Supplementary Fig. S7c and S7d), bulk RNA-seq data were obtained from ArrayExpress (E-MTAB-4748, https://www.ebi.ac.uk/biostudies/arrayexpress), and only female iPSC lines were extracted for downstream analysis. All analyses were performed in R. To assess the relationship between *XIST* and *XACT* expression, Pearson correlation coefficients were calculated, and correlation plots were generated. Statistical significance of the correlation was evaluated using the Pearson correlation test.

For comparative analyses, iPSC lines were stratified into two groups based on log2-transformed TPM values: *XACT*-high (*XACT* log2TPM ≥ 1.5 and XIST log2TPM ≤ 1) and *XACT*-low (*XACT* log2TPM ≤ 1 and XIST log2TPM ≥ 2). The expression levels of naïve pluripotency- and differentiation-related genes were then compared between these groups.

### Human axioloid formation from hPSCs

The derivation of axioloid-like structures was performed as previously described ^41^. Briefly, hPSCs were seeded onto 6-well plates coated with iMatrix-511 silk (Nippi, Kusatsu, Shiga, Japan) and cultured for 5 days until they reached approximately 60% confluency. Axioloid induction was carried out using AK02N medium without component C (ReproCELL, Yokohama, Kanagawa, Japan). To initiate differentiation, hPSCs were treated for 24 hours with basic FGF (20 ng/mL) (STEMCELL Technologies, Vancouver, Canada) and the WNT agonist CHIR99021 (5 μM) (Tocris Biosciences, *Bristol, UK*) to promote mesoderm and primitive streak fates. After this initial treatment, cells were dissociated using Accutase (Thermo Fisher Scientific, Waltham, MA, USA) and transferred to 96-well U-bottom low-attachment plates (Corning, Inc, Arizona, USA) at a density of 1,000 cells per well. Aggregation was carried out in 50 μL of AK02N-C-based medium supplemented with CHIR99021 (5 μM), basic FGF (20 ng/mL), the TGFβ inhibitor SB431542 (10 μM) (STEMCELL Technologies, Vancouver, Canada), and Y27632 (10 μM) for 24 hours. After 72 hours of aggregation, individual axioloids were transferred to 96-well flat-bottom low-attachment plates (Corning, Inc, Arizona, USA) pre-treated with BSA. They were embedded in 80 μL of AK02N-C medium containing 10% growth factor-reduced Matrigel (Corning, Inc, Arizona, USA) and cultured at 37°C with 5% CO□ for an additional 24 to 48 hours. Depending on the experimental conditions, retinoic acid (RA, 100 nM) (STEMCELL Technologies, Vancouver, Canada) was included in the Matrigel-containing medium during the 96 low attach flat plate (AGC TECHNO GLASS Co.,Ltd., Shizuoka, Japan) embedding phase. For *XACT*^VPR^-3 and *XACT*^VPR^-6 lines, cells were pretreated with Dox for 144 hours prior to axioloid induction.

### Axioloids Immunofluorescence Combined with RNA-FISH

Axioloid samples were washed once with PBS containing 0.1% PVA (Sigma-Aldrich., St. Louis*, USA*) (PBS-PVA) and fixed in 4% paraformaldehyde (PFA) at room temperature for 20 minutes. Following fixation, the samples were washed three times with PBS-PVA. Permeabilization was performed using 0.2% Triton X-100(Sigma-Aldrich., St. Louis*, USA*) in PBS for 20 minutes at room temperature, followed by blocking with 1.5% BSA and RNaseOUT in PBS for 1 hour at room temperature. For immunostaining, the samples were incubated with primary antibodies diluted in 1.5% BSA with RNaseOUT for 1 hour at room temperature. The primary antibodies used included: rabbit anti-NANOG (1:100, Abcam), mouse anti-OCT4 (1:300, Santa Cruz), goat anti-SOX2 (1:100, R&D Systems), rabbit anti-CDX2 (1:100, R&D Systems), rabbit anti-TBXT (1:100, Cell Signaling), goat anti-GATA6 (1:100, R&D Systems), and goat anti-TBX6 (1:100, R&D Systems). The samples were then washed three times with PBS-PVA and incubated with secondary antibodies (1:500 dilution) in PBS for 1 hour at room temperature. After washing the samples three times with PBS-PVA, they were stained with DAPI for 10 minutes at room temperature. Secondary antibodies used included: Goat anti-rabbit AlexaFluor 488, Goat anti-mouse AlexaFluor 633, donkey anti-goat AlexaFluor 633, and donkey anti-goat AlexaFluor 488 (all from Thermo Fisher Scientific). After DAPI staining, the specimens were post-fixed with 4% PFA for 20 minutes, washed three times with PBS-PVA, and placed on glass slides. The slides were coated with 70% ethanol, followed by 100% ethanol, and left to dry at room temperature for 5 minutes before the RNA-FISH assay.

For RNA-FISH, 4 μL of a 1:1 mixture of hybridization buffer and *XACT*-cy3 probe was applied, covered with Parafilm (Bemis Company, I*nc.,* Wisconsin, USA), and incubated overnight at 37°C in a humidified chamber. The following day, the samples were washed twice in formamide at 45°C for 20 minutes, followed by two washes in 0.05% Tween-20 at 45°C for 20 minutes. The samples were then washed with 70% and 100% ethanol, left to dry for 5 minutes at room temperature, and mounted with 5 μL of DAPI-free ProLong Glass Antifade Mountant (Thermo Fisher Scientific, Waltham, MA, USA). Coverslips were applied, and the specimens were sealed. Fluorescence imaging was performed using a Carl Zeiss LSM880 microscope.

### Quantification of Axioloids

Raw image files were imported into Imaris software (v. x64 9.0.2, Oxford Instruments). Total cell numbers were detected using the SPOT function based on DAPI staining. The threshold parameters were adjusted to accurately detect cells. Each stained cell was manually measured. The proportion of stained cells was calculated based on these measurements. Each sample had n=5, and graphs were generated using R.

## Author Contributions

K.I. and A.F. conceived the study and designed the experiments. K.I. performed the majority of experiments and analyses, with assistance from N.M., A.S., and Natsumi K. K.I., Y.I., I.H., K.Y., Natsuhiko.K., and A.F. carried out single-cell experiments and data analysis. M.Y., A.U., H.I., and H.A. provided iPSC lines and contributed to data interpretation. K.I. and A.F. wrote the manuscript with input from all authors. A.F. supervised the project.

## Acknowledgements

We thank the members of the Support Center for Medical Research and Education at Tokai University School of Medicine for their assistance with various experiments. This study was supported by an institutional grant to K.I. (2022 Tokai University School of Medicine Research Aid, 2023 Tokai University School of Medicine Research Aid) and by JSPS KAKENHI Grant Number 23H00436 to A.F.

## Supplementary Figure legends

**Supplementary Figure S1. Generation of an inducible *XACT* line and hyper-elevation of NANOG and OCT4 in *XACT*^VPR-6 cells.** (a) Schematic of the dCas9-VPR–mediated *XACT* activation system. A PiggyBac vector system was used to express Dox-inducible dCas9-VPR and gRNAs targeting the *XACT* locus. Following transfection, hPSCs were selected with hygromycin and puromycin. (b) RNA-FISH analysis of *XACT* in WT, *XACT*^Δ/Y^-11, and *XACT*^VPR^-6 cells. Experimental scheme as in Fig. 1b. Scale bar, 20 µm. (c) qPCR analysis of *XACT* induction in *XACT*^VPR^-6 cells. Each dot represents a biological replicate. *P < 0.05 (Student’s t-test). (d) Bright-field images of *XACT*^Δ/Y^-11 and *XACT*^VPR^-6 cells cultured in SF or Un.diff. conditions. Scale bars, 100 µm. (e) Crystal violet assay of cell growth in Un.diff. conditions. Cells were seeded at 1 × 10^5^ and stained at 48 and 144 h post-Dox. Each dot represents a biological replicate; bar graphs indicate mean values. *P < 0.05; **P < 0.01 (Student’s t-test). Scale bar, 5 mm. (f) qPCR analysis of pluripotency and differentiation markers in *XACT*^VPR^-6 cells. Sampling scheme as in Fig. 1f. *P < 0.05; **P < 0.01 (Student’s t-test). (g) WB analysis of pluripotency factors in *XACT*^VPR^-6 cells. GAPDH served as a loading control. Representative blots are shown (uncropped images and biological replicates in Supplementary Fig. S10).

**Supplementary Figure S2. Heterogeneous and reversible OCT4 and NANOG expression in *XACT*^VPR-6 cells.** (a) IF analysis of pluripotency factors in *XACT*^VPR^-6 cells. NANOG was imaged at different exposures to assess heterogeneity. DAPI stained nuclei. Scale bar, 100 µm. (b) IF-FISH analysis of NANOG/OCT4/ *XACT* in *XACT*^VPR^-6 cells cultured in SF or Un.diff. conditions. Scale bar, 20 μm. (c–d) Correlation of *XACT* expression with NANOG or OCT4 (c), and between NANOG and OCT4 (d), at single-cell resolution. Each dot represents one cell. Pearson correlation; significance by two-tailed t-test. (e) Propagation assay of *XACT*^VPR^-6 cells cultured in Un.diff. conditions and passaged with TrypLE. Scale bar, 5 mm. (f) IF-FISH analysis of *XACT* and pluripotency factors. Scale bar, 20 µm. (g) Crystal violet staining of cell growth. Scale bar, 100 µm. (h) Reversibility assay: IF-FISH of *XACT* and pluripotency markers with or without Dox. Scale bar, 50 µm.

**Supplementary Figure S3. Continuous *XACT* expression impairs directed differentiation in *XACT*^VPR^-6 cells.** (a) Crystal violet staining of cells cultured in lineage-specific differentiation media. Scale bar, 5 mm. (b) IF-FISH analysis of pluripotency factors and lineage markers (ectoderm: PAX6; mesoderm: TBXT). Endoderm differentiation was not achieved due to insufficient cell numbers. DAPI stained nuclei. Scale bar, 50 µm. (c) Quantification of marker-positive cells. At least 115 cells were analyzed. (d) Quantification of *XACT* expression relative to each marker.

**Supplementary Figure S4. Molecular features of *XACT*-altered cells.** (a) Cell cycle analysis from scRNA-seq data. (b) Growth status of *XACT*^VPR^-3 cells cultured in SF, assessed by DAPI staining. Scale bar, 2 cm. (c) UMAP showing cluster identities by genotype. (d) Marker expression status: (i) VIM (differentiation) and (ii) DPPA3 (naïve). (e) GO analysis of DEGs in *XACT*^VPR+^ cells. Adjusted p-values were calculated using the Benjamini–Hochberg method (FDR).

**Supplementary Figure S5. *XACT* is highly expressed in seminoma.** (a) *XACT* expression across cancer types from UALCAN (PMID: 35078134). (b) UMAP visualization of naïve hPSCs, derivatives, and seminoma. (i) Cell line batch. (ii) Clusters. (c) Expression of *XACT/*NANOG/GDF3. (d) Cell clusters used as reference for prediction analysis in Fig. 4f.

**Supplementary Figure S6. Sequence analysis of 3′ UTRs in *XACT*^VPR+ cells.** (a) Full RT-PCR gel images from 3’ RACE. Amplicons (dotted lines) were sequenced and analyzed by BLAST (https://blast.ncbi.nlm.nih.gov/Blast.cgi). (b) Genomic PCR of OCT4 and NANOG 3’ UTRs. Primer sites, full gels, and sequence results (dotted lines) are shown. Sanger sequencing was performed in *XACT*^VPR^-3 cells.

**Supplementary Figure S7. Female hPSCs with *XACT*^high/XIST^low expression upregulate naïve pluripotency markers.** (a) *XIST/XACT* RNA-FISH in adsc-iPSCs. Scale bar, 20 µm. (b) Bulk RNA-seq analysis (GSE218950). Two biological replicates of adsc-iPSCs in SF. Bars show fold change relative to average TPM. (c) *XACT* and *XIST* expression in female HipSci iPSCs (n = 106). Pearson correlation and P values are shown. Each dot = one line; 49 highlighted lines (*XACT*^high/*XIST*^low or *XACT*^low/*XIST*^high) were used in (d). (d) Expression of core pluripotency, naïve, and differentiation genes. *P < 0.05; **P < 0.01 (Student’s t-test).

**Supplementary Figure S8. Super-resolution DNA-FISH of *XACT*, *GDF3*, *DPPA3*, and *NANOG* loci.** (a) Genomic regions and probe positions. Assays were performed under SF with or without Dox for 144 h. (b) DNA-FISH images and quantification of *XACT* signal length. *P < 0.01 (Student’s t-test). Scale bar, 5 µm.

**Supplementary Figure S9. Axioloid modeling using *XACT***^Δ**/Y**^**-11 and *XACT*^VPR^-6 cells.** (a) Bright-field images of axioloid morphology. Scale bar, 100 µm. (b) Quantification of axioloid diameter at 48 h. *P < 0.05; **P < 0.01 Statistical significance was determined by one-way ANOVA with Tukey’s multiple-comparison test. (c–d) IF-FISH of *XACT* and lineage markers. (c) IF images: pluripotency (NANOG, OCT4, SOX2), trophectoderm (CDX2), primitive endoderm (GATA6), and mesoderm (TBXT, TBX6). Percentages of marker-positive cells are shown. Scale bar, 20 µm. (d) Representative *XACT* RNA-FISH. Scale bar, 20 µm.

**Supplementary Figure S10. Uncropped western blot images.**

