## Supplementary figures and images for "Upregulation of the lncRNA *XACT* sustains pluripotency, blocks lineage specification, and drives germ cell tumor-like transcriptional programs in human pluripotent stem cells"

### supplemental Figure

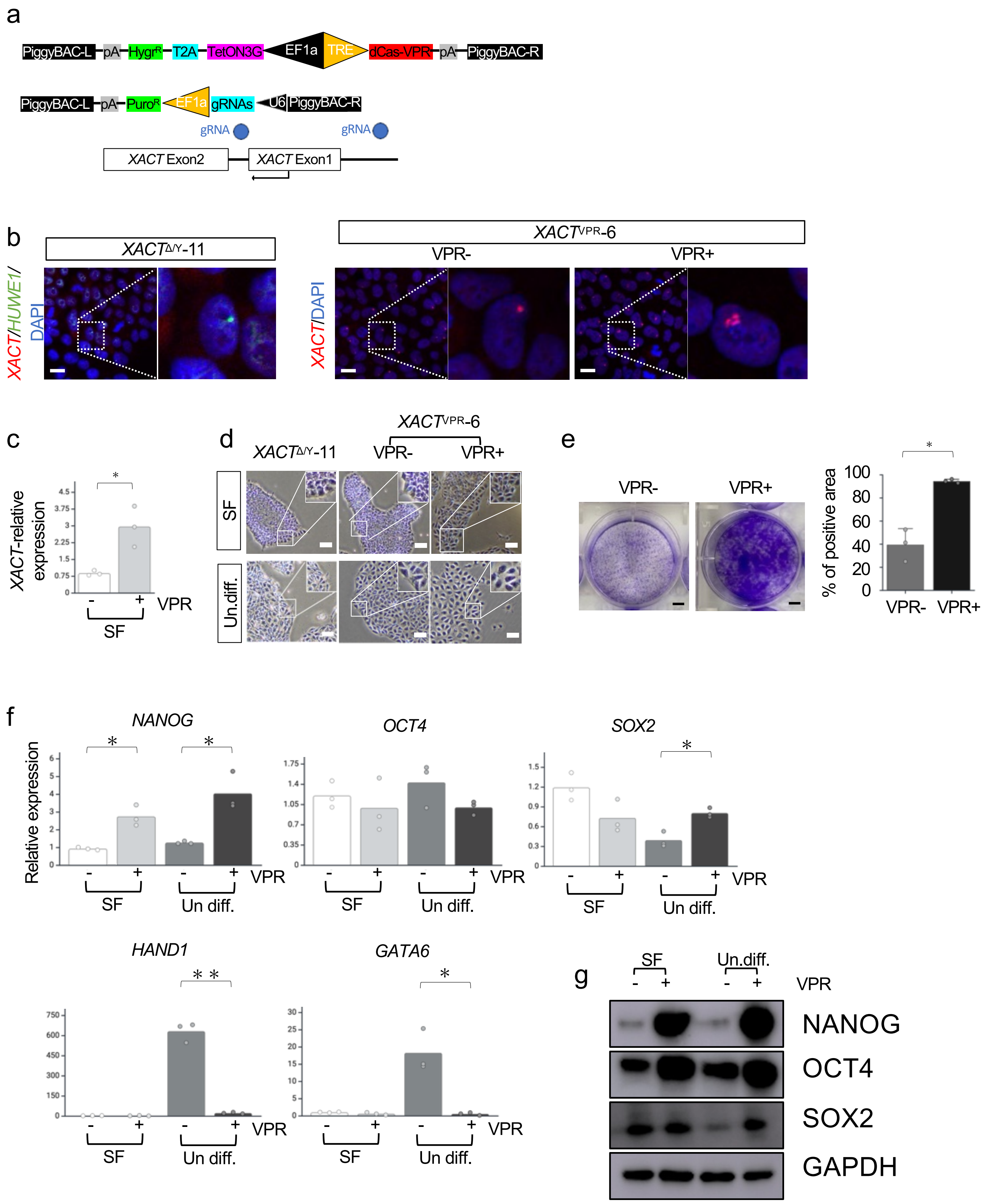

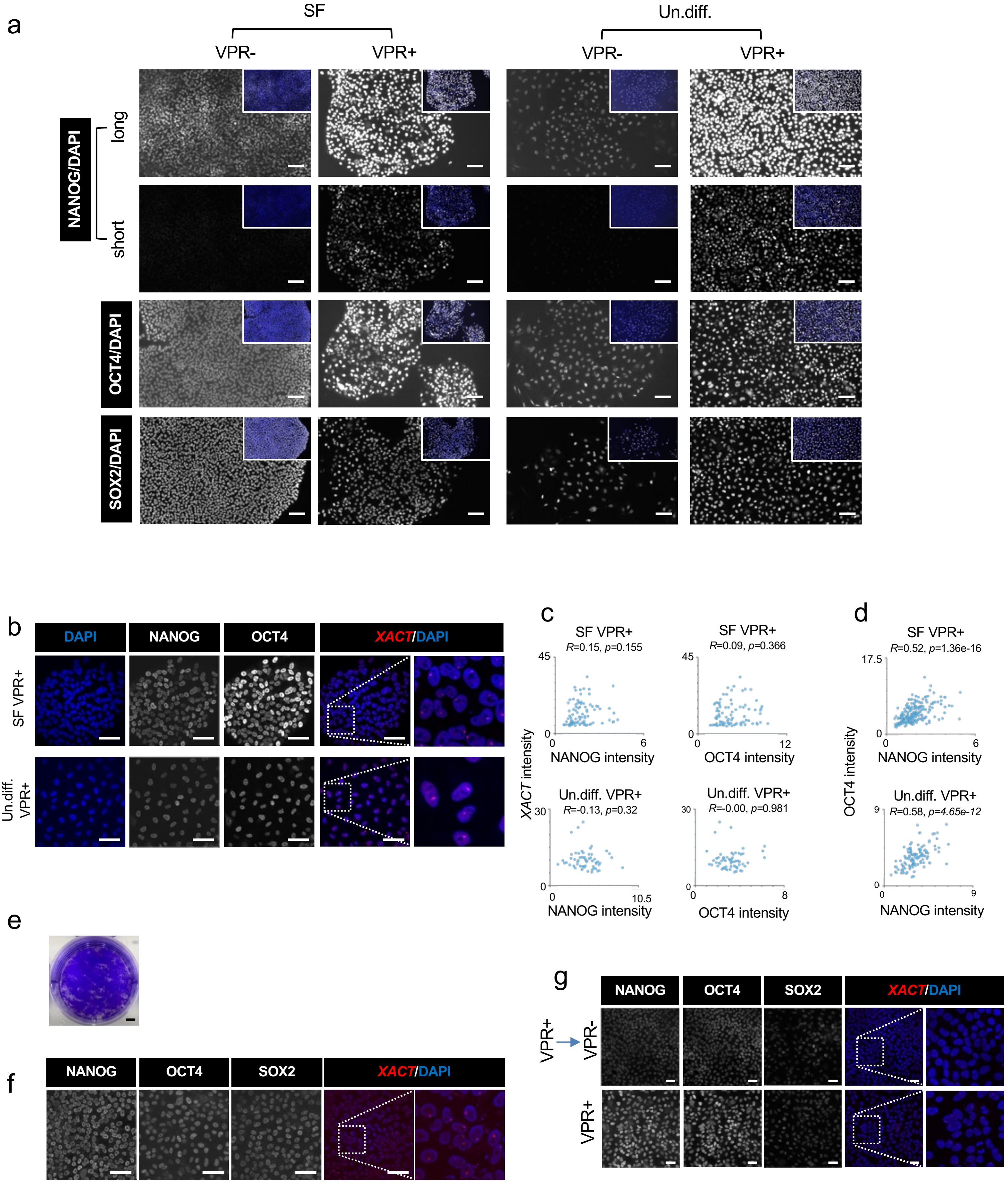

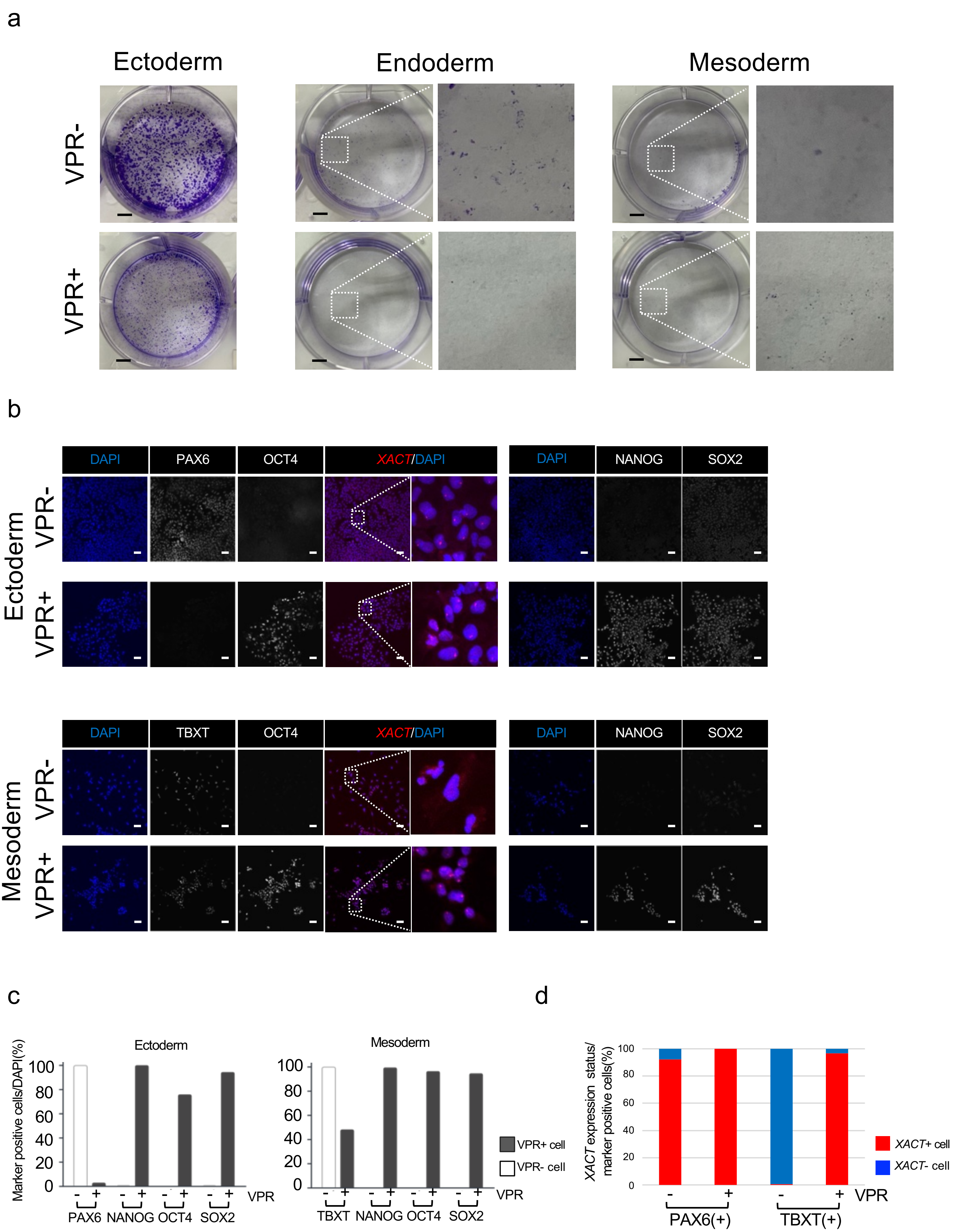

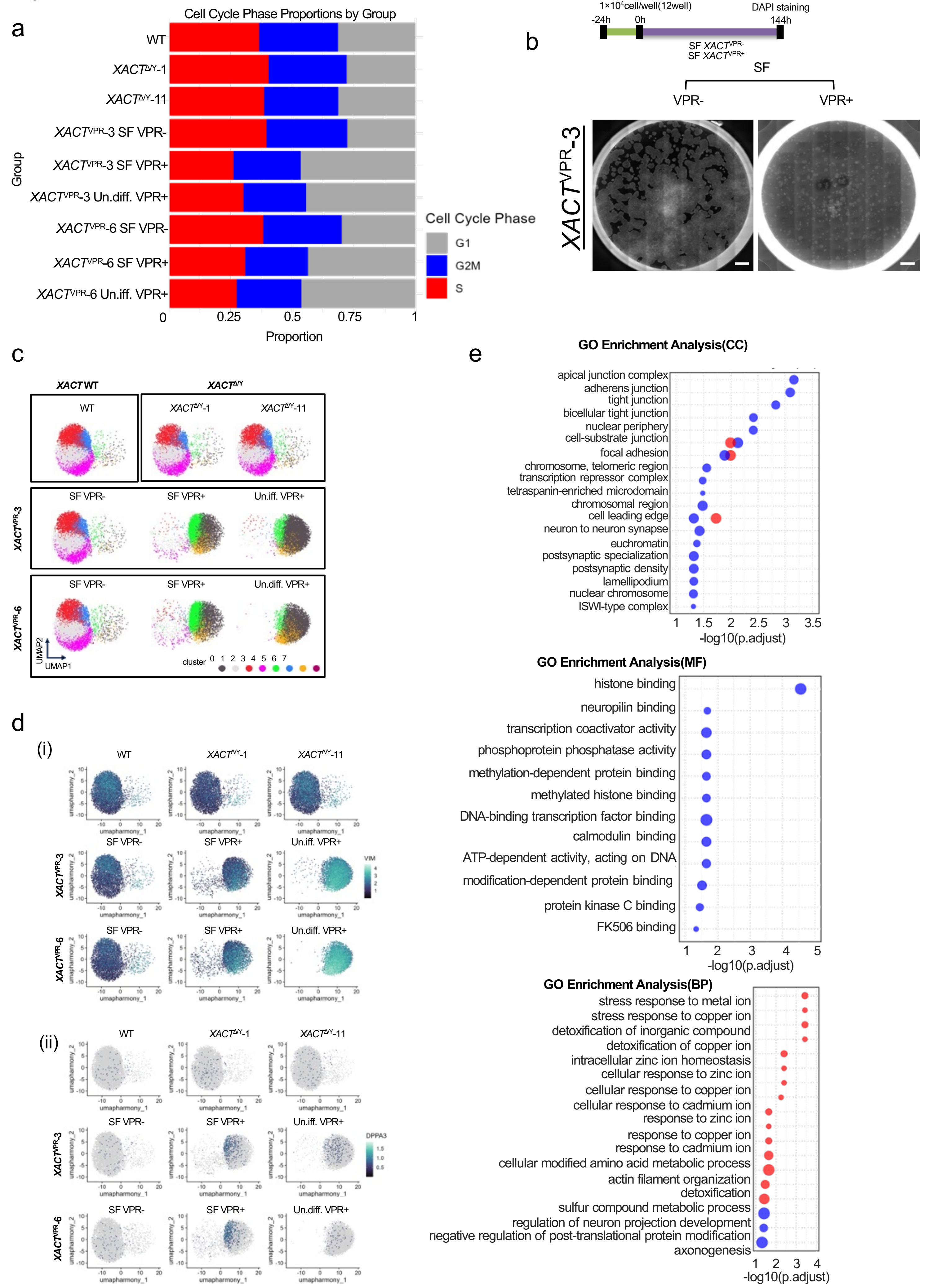

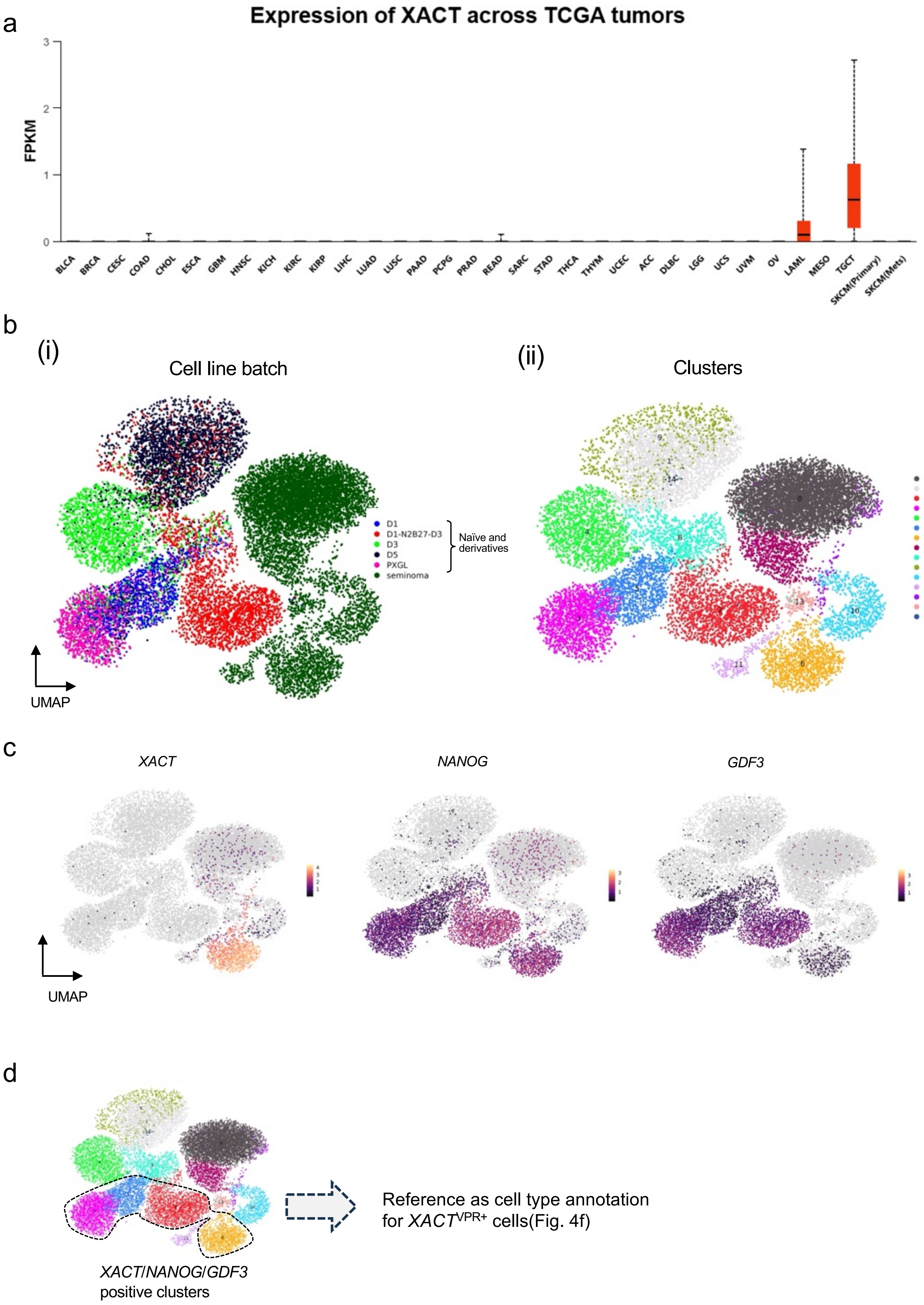

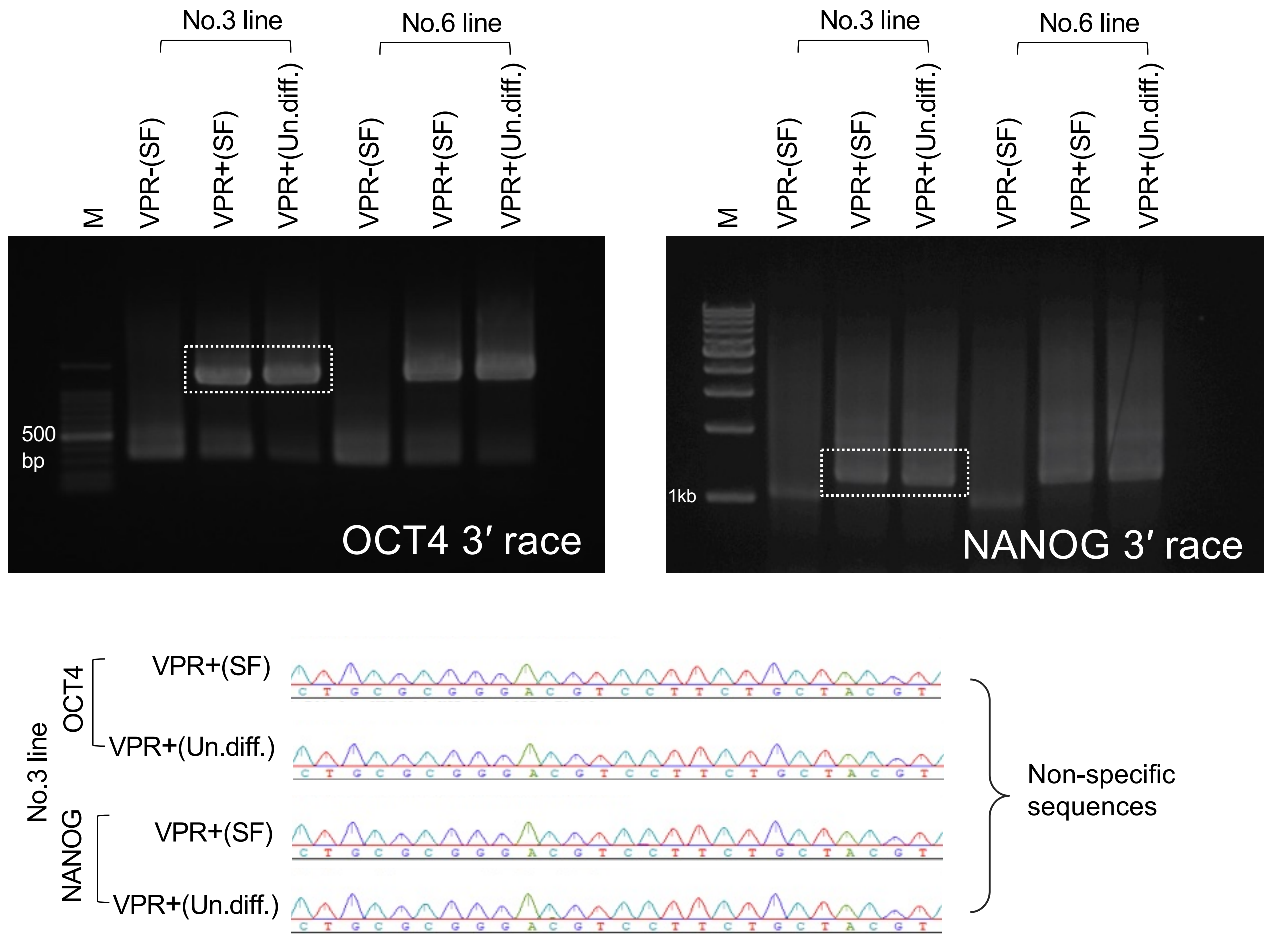

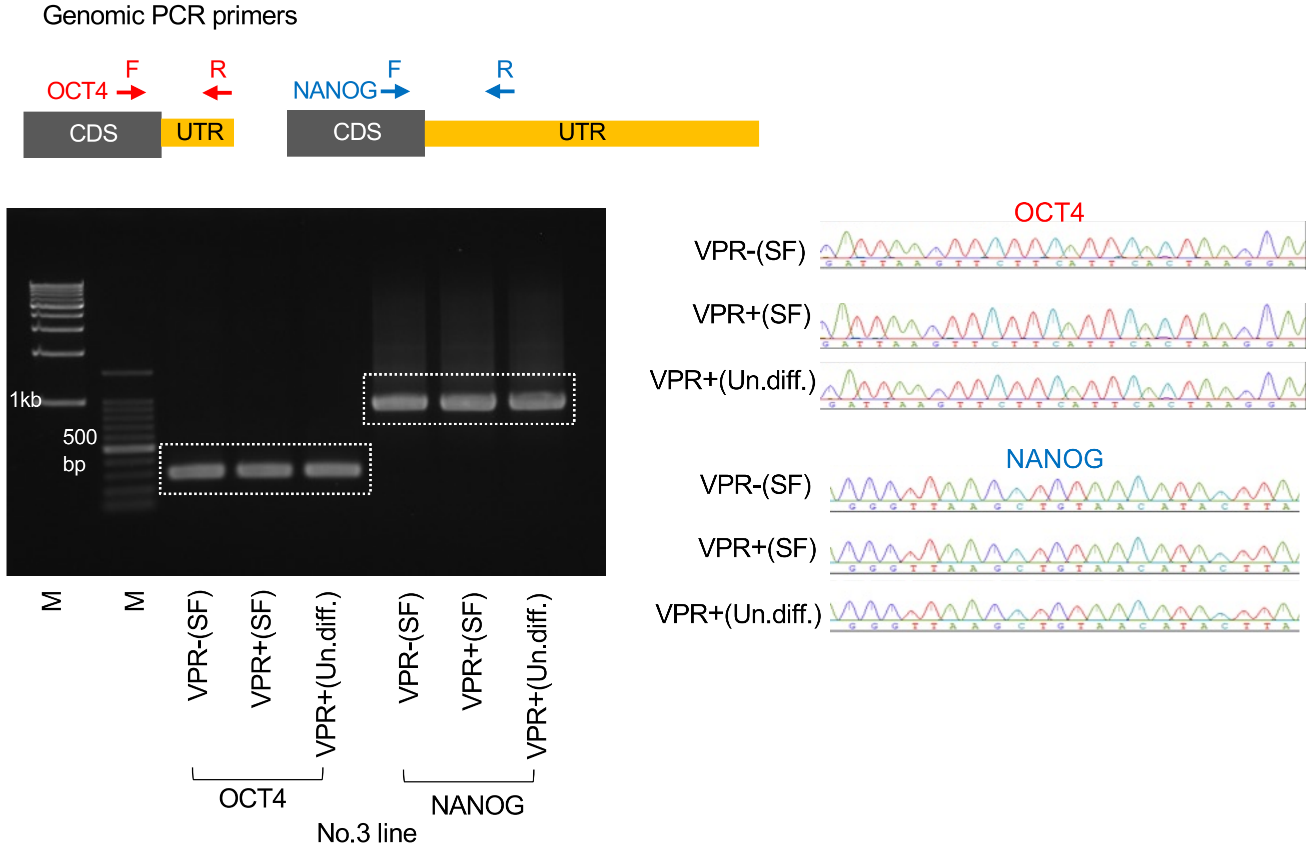

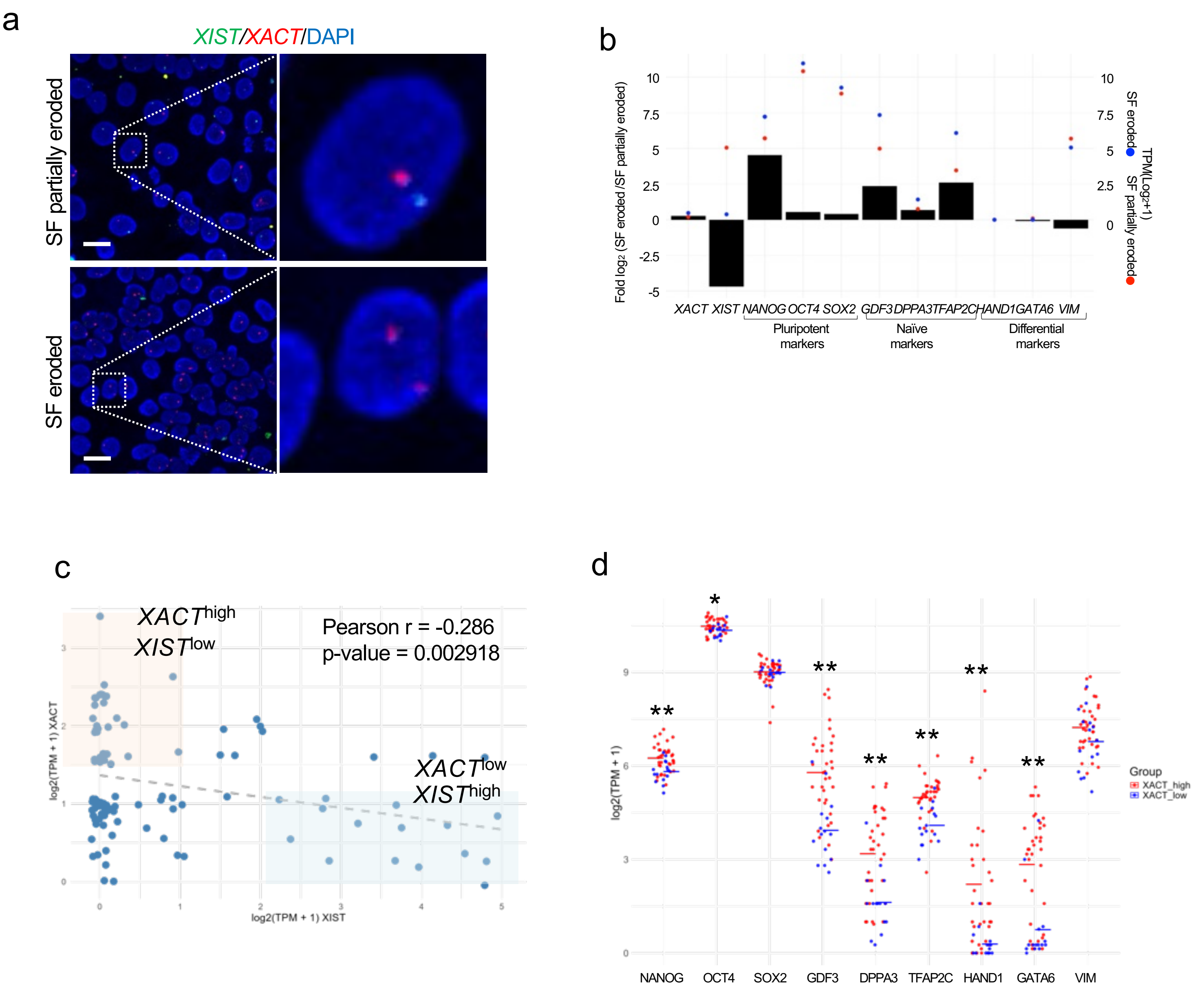

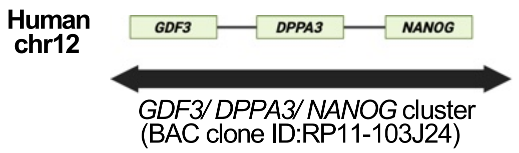

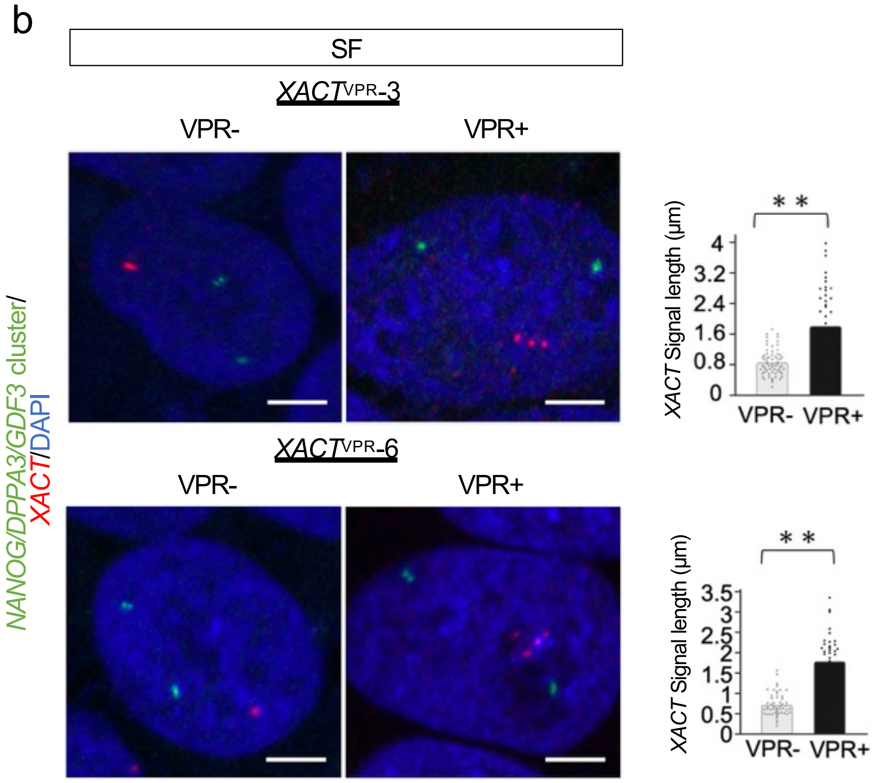

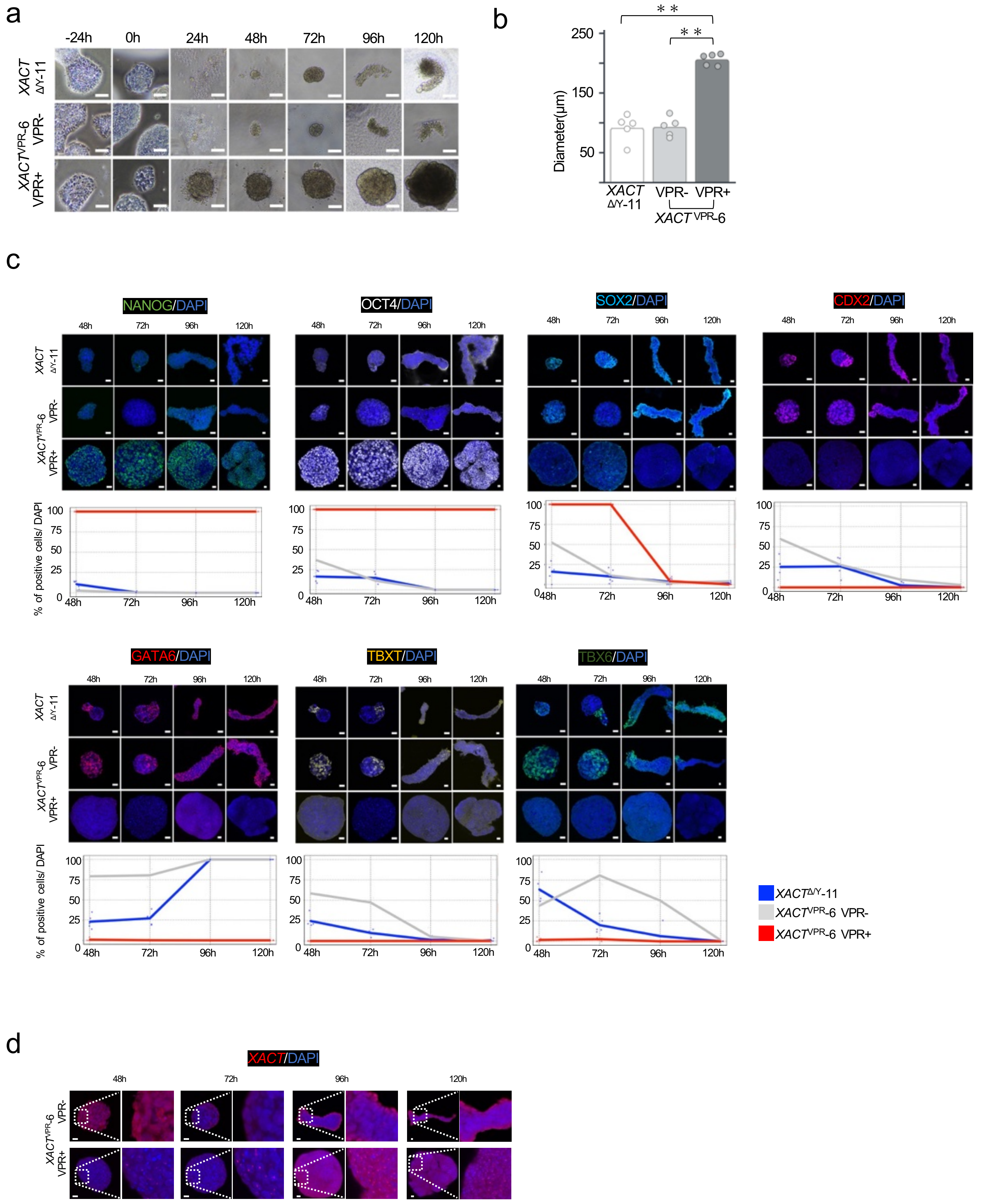

**Fig. S10**

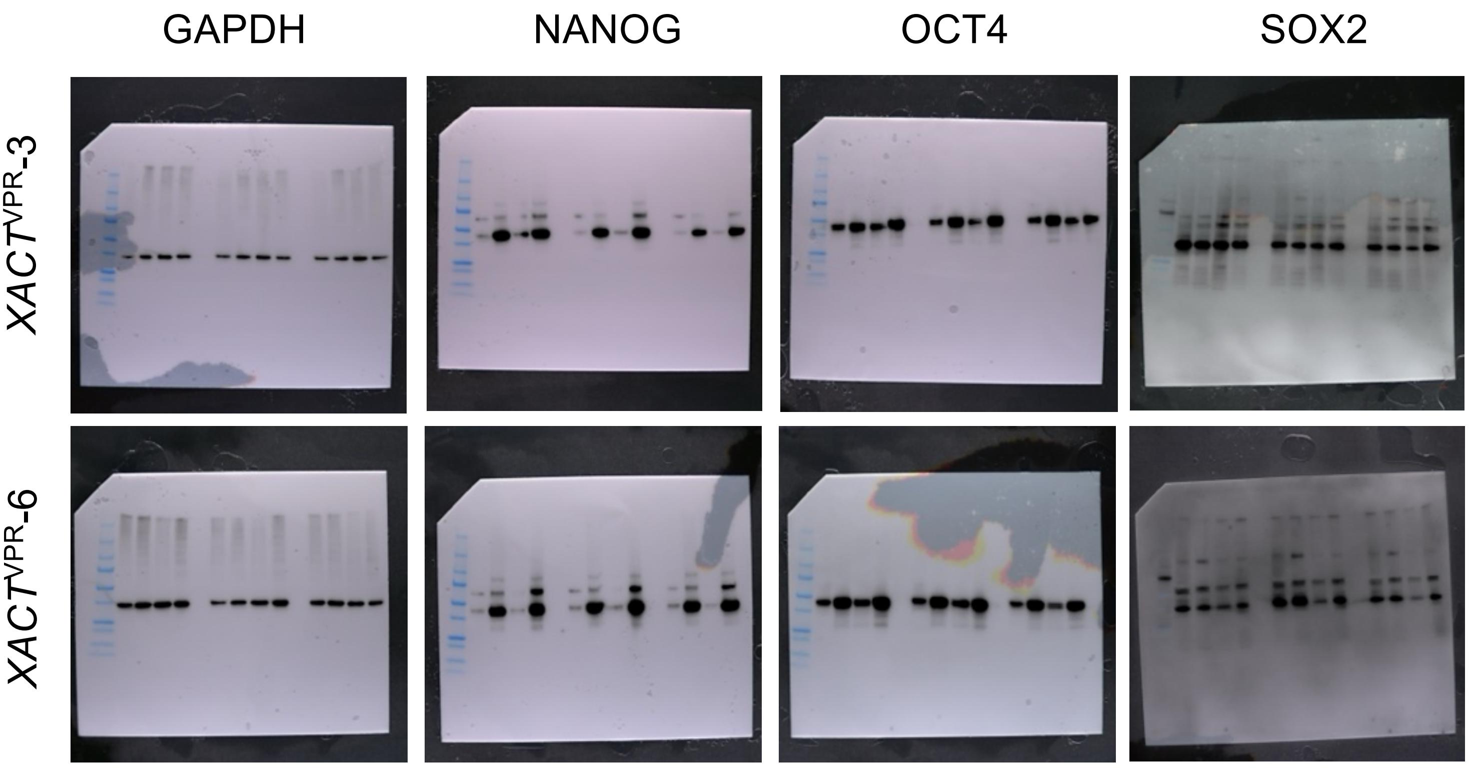
